# Natural killer cell regulation of breast cancer stem cells mediates metastatic dormancy

**DOI:** 10.1101/2023.10.02.560493

**Authors:** Grace G. Bushnell, Deeksha Sharma, Henry C. Wilmot, Michelle Zheng, Toluwaleke D. Fashina, Chloe M. Hutchens, Samuel Osipov, Max S. Wicha

**Affiliations:** University of Michigan

## Abstract

Breast cancer patients with estrogen receptor positive tumors face a constant risk of disease recurrence for the remainder of their lives. Dormant tumor cells residing in tissues such as the bone marrow may generate clinically significant metastases many years after initial diagnosis. Previous studies suggest that dormant cells display “stem like” properties (CSCs), which may be regulated by the immune system. Although many studies have examined tumor cell intrinsic characteristics of dormancy, the role of the immune system in controlling dormancy and its escape is not well understood. This scientific gap is due, in part, to a lack of immunocompetent mouse models of breast cancer dormancy with many studies involving human xenografts in immunodeficient mice. To overcome this limitation, we studied dormancy in immunocompetent, syngeneic mouse breast cancer models. We find that PyMT, Met-1 and D2.0R cell lines contain CSCs that display both short- and long-term metastatic dormancy *in vivo*, which is dependent on the host immune system. Natural killer cells were key for the metastatic dormancy phenotype observed for D2.0R and the role of NK cells in regulating CSCs was further investigated. Quiescent D2.0R CSC are resistant to NK cytotoxicity, while proliferative D2.0R CSC were sensitive to NK cytotoxicity both *in vitro* and *in vivo*. This resistance was mediated, in part, by the expression of Bach1 and Sox2 transcription factors. NK killing was enhanced by the STING agonist MSA-2. Collectively, our findings demonstrate the important role of immune regulation of breast tumor dormancy and highlight the importance of utilizing immunocompetent models to study this phenomenon.

## Introduction

Metastatic dormancy is a significant clinical problem for the 3.8 million women in the United States with breast cancer (*1*). Most women will have early-stage diagnosis and receive therapy that is curative on the scale of 5-10 years. However, many of these patients will recur with metastatic disease 10 years or more after initial diagnosis (*2*). The risk factors for recurrence are unclear, but evidence suggests a role for immune system function (*3*). Most mouse models of metastatic dormancy have utilized human tumor cells inoculated into immunocompromised mice (*3*) limiting their ability to uncover key immune components and thus their clinical relevance.

Natural killer (NK) cells are a major component of the immune system that has been implicated in controlling both metastasis (*4*) and metastatic dormancy (*5, 6*). NK cells are innate immune cells that can distinguish normal cells from virus infected and transformed cancer cells without requiring antigen specificity like T cells or B cells. NK cells receive signals from both activating and inhibitory receptors on target cells and integrate these signals in the context of the tissue microenvironment to regulate cellular killing. Activating receptors such as NKG2D, natural cytotoxicity receptors, DNAM, and CD16, among others (*7*) recognize “induced self” signals on target cells upregulated during viral infection, inflammation, or cellular stress. Inhibitory receptors such as killer immunoglobulin receptors (KIRs), NKG2A, and others, recognize MHC class I molecules present on the surface of target cells. Tumor cells and virally infected cells frequently downregulate MHC class I, allowing them to escape recognition of mutated or viral antigens by T cells. As a result, a compensatory mechanism for NK cell regulation of these cells with “missing self” signals evolved (*7*). Similarly, clinically relevant tumor in a human or a mouse has escaped this regulation through modulation of these receptors or inhibition of NK cell function in the tumor or metastatic microenvironment (*8*).

The paucity of immunocompetent mouse models has hindered research into the role of the immune system in regulating tumor dormancy. To address this, we evaluated murine breast cancer cell lines for their short-term and long-term dormancy in fully immunologically competent mouse models and found that each are regulated by different components of the immune system. We focused on the D2.0R model which exhibits NK cell dependent metastatic dormancy via quiescent D2.0R cancer stem cells (CSC) evading NK cell-mediated immunity through downregulation of activating NK ligands, STING and STING targets, and upregulation of MHC partially controlled by Bach1 and Sox2 expression.

## Results

### Short- and long-term dormancy model development

We chose five murine mammary adenocarcinoma cell lines (PyMT, Met-1, D2.0R, D2A1, and E0771) to investigate for long-term dormancy following intracardiac inoculation. We injected 100,000 PyMT, Met-1, D2.0R, D2A1, or E0771 murine mammary carcinoma cells into the left ventricle of syngeneic mice and monitored mouse health over time (**Fig. 1a**). Our experiments revealed that these cell models fit two phenotypes: cell lines that form metastasis very quickly and cell lines that exhibit an extended period of metastatic dormancy (greater than one hundred days) (**Fig. 1b**).

**Figure 1:**
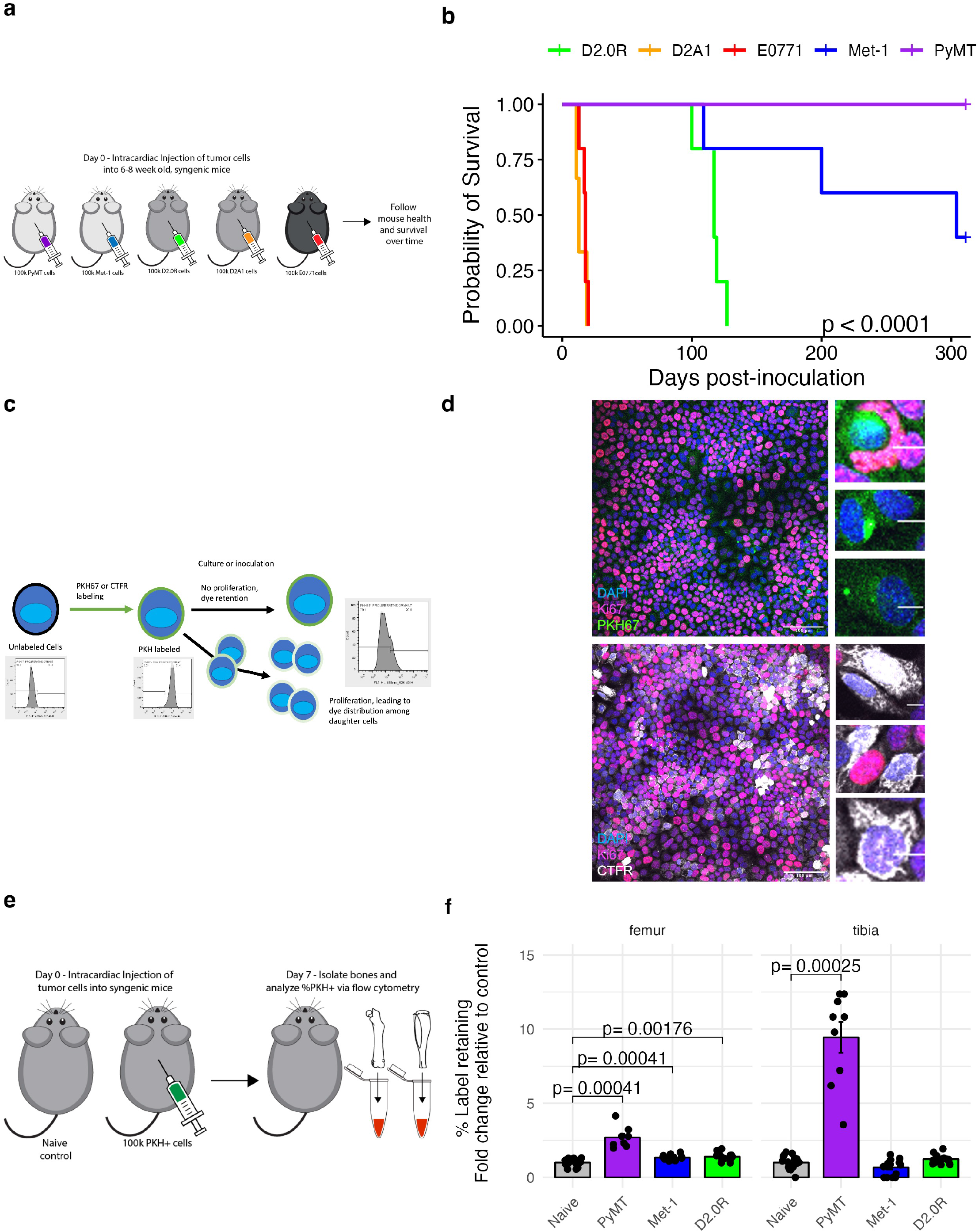
PyMT, Met-1, and D2.0R murine mammary carcinoma cell lines exhibit long-term metastatic dormancy and short-term quiescence in bone marrow. (**a**) Schematic for evaluation of long-term dormancy in vivo. (**b**) Evaluation of survival for mice injected with 100,000 PyMT, Met-1, D2.0R, D2A1, or E0771 cells. (**c**) Schematic for fluorescent label retention as a readout for quiescent cells. (**d**) Representative fluorescence micrographs for D2.0R cells labeled with PKH67 (top) or CTFR (bottom) and allowed to proliferate for 7 days in vitro before antibody labeling with Ki-67 to determine the association of label-retention and Ki-67 proliferation marker. (**e**) Schematic for fluorescent label-retention as a readout for quiescent cells in short-term dormancy assay in vivo. (**f**) Quantification of label retaining cells in femurs or tibias of mice injected with PyMT, Met-1, or D2.0R cells normalized to the naïve control. P-values were calculated using t test (for normally distributed data) or Wilcoxon test with Bonferroni correction for multiple comparisons when appropriate (* p < 0.05, ** p < 0.01, *** p < 0.001, **** p<0.0001). Survival curves were compared via log-rank test calculation of p-value.

D2A1 and E0771 belong to the first group with mice exhibiting extensive metastases within 30 days. PyMT, Met-1, and D2.0R belong to the second group with mice surviving past one hundred days.

We next developed a model of short-term dormancy of breast cancer cells in bone marrow following intracardiac injection in immunocompetent syngeneic mice. We utilized fluorescent label retention (*9*) (PKH67 or cell trace far red, CTFR) to label non-proliferating tumor cells both *in vitro* and *in vivo* (**Fig. 1b**). Cells that retain the label are quiescent label-retaining cells (LRC) while cells that do not retain the label (non-LRC) are proliferative, as confirmed by Ki-67 staining (**Fig. 1c**). Another advantage of this labelling technique is that unlike genetically introduced tags such as RFP, which elicit an immune response (**Fig. S1a**), these dyes are non-immunogenic (**Fig. S1b-c**). We used these dyes to label PyMT, Met-1, and D2.0R murine mammary carcinoma cell lines and injected these cells into syngeneic mice (FVB/N for PyMT and Met-1 and Balb/c for D2.0R) via intracardiac injection to allow the cells to distribute throughout the circulation (**Fig. 1d**). LRC were detectable in the femurs of mice injected with PyMT, Met-1, and D2.0R cells and tibias of mice injected with PyMT cells (**Fig. 1e**). This short-term model of dormancy indicates that non-proliferative tumor cells lodge in bone marrow and can be detected after one week. Taken together these data indicate that PyMT, Met-1, and D2.0R exhibit both short- and long-term dormancy, while D2A1 and E0771 do not. As a result, we chose to further investigate the PyMT, Met-1, and D2.0R cell lines.

### Evaluation of murine breast cancer stem cell populations in dormant cell lines

After identifying PyMT, Met-1, and D2.0R as murine mammary carcinoma cell lines that display long-term metastatic dormancy in vivo, we investigated the CSC populations in these cell lines (D2.0R **Fig. 2**, PyMT, Met-1 **Fig. S2**) by staining for ALDH, Sca-1, and CD90 (see **Fig. S3** for gating scheme) which are well-established markers of CSC in other models of human and murine breast cancer (*10*). We compared 2D (2% FBS and tissue culture treated flasks) and sphere formation conditions for each cell line (**Fig. 2a, Fig. S2a**). Mammosphere culture enriches for a proliferative stem-like phenotype (*11*), while low serum 2D culture enriches for a quiescent stem-like population (*12*). The enrichment of ALDH^+^ and Sca-1^+^CD90^-,^ and the depletion of Sca-1^-^CD90^+^ in a more proliferative sphere culture relative to quiescent 2D cultures, suggests that the ALDH^+^ and Sca-1^+^CD90^-^ CSC are more proliferative, while the Sca-1^-^CD90^+^ CSC are more quiescent. To test this hypothesis, we performed combined staining of the CSC panel with CTFR labeling (**Fig. 2b**). We compared the expression of CSC markers in 2D culture for all live cells, LRC, or non-LRC to determine whether these populations are quiescent (LRC) or proliferative (non-LRC). The ALDH^+^ and Sca-1^+^CD90^-^ populations are significantly reduced in LRC relative to live, confirming that these populations are proliferative while the Sca-1^-^CD90^+^ population was significantly enriched in the LRC relative to live, indicating that this population is quiescent.

**Figure 2:**
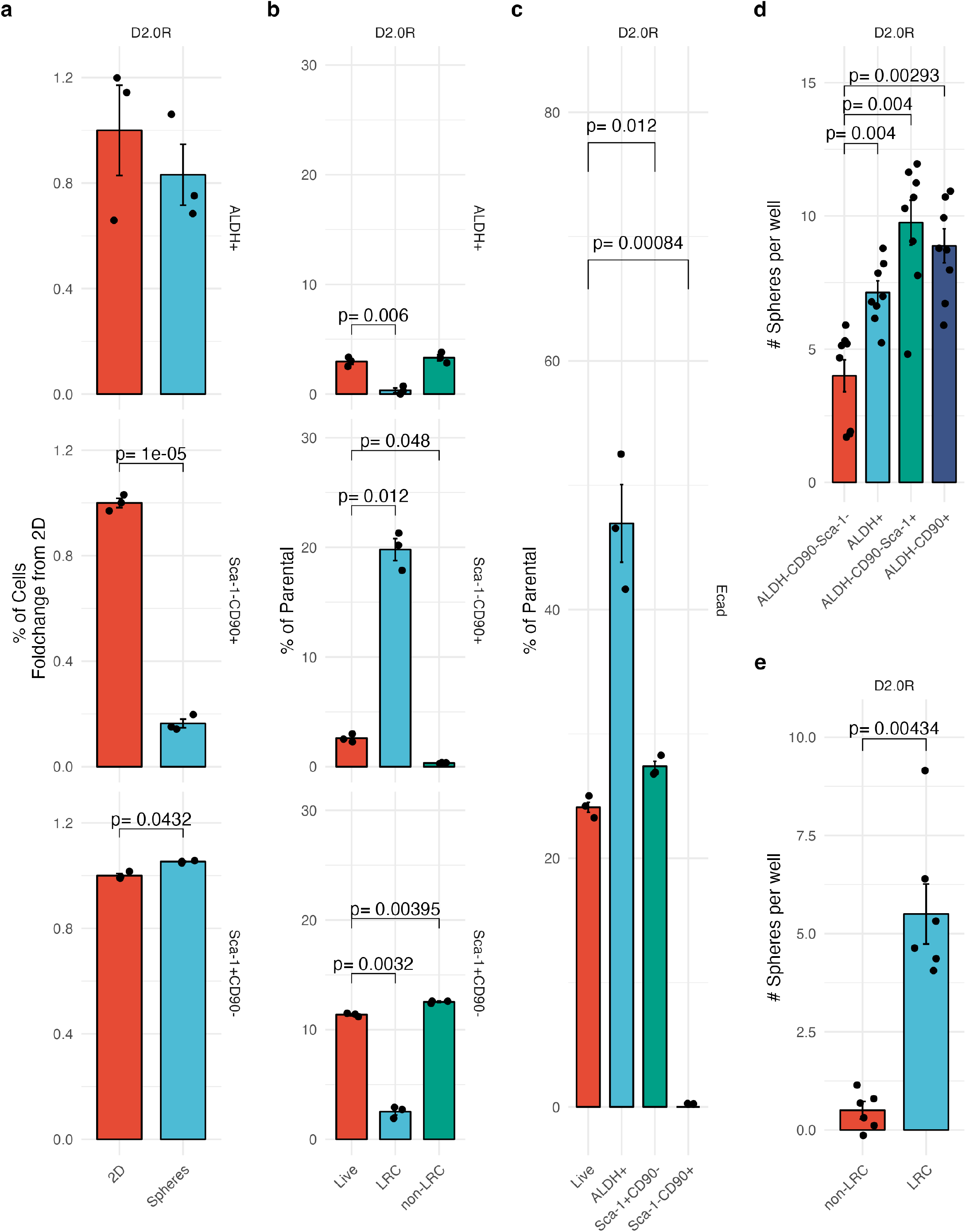
Murine cell lines contain an epithelial, proliferative cancer stem cell and a mesenchymal, quiescent cancer stem cell population. (**a**) Flow cytometry evaluation of cancer stem cell markers ALDH (top), Sca-1^-^ CD90^+^ (middle) and Sca-1^+^ CD90-(bottom) for D2.0R cell line in 2D (2% FBS tissue-culture treated flasks) and 3D (mammosphere media ultra-low attachment flasks) reported as fold change from 2D. (**b**) Flow cytometry evaluation of cancer stem cell markers ALDH (top), Sca-1^-^ CD90^+^ (middle) and Sca-1^+^ CD90-(bottom) for D2.0R cell line in 2D (2% FBS tissue-culture treated flasks) as a percentage of parental population (Live, LRC or non-LRC). (**c**) Flow cytometry evaluation of E-cadherin (Ecad) as a percentage of parental population (Live, ALDH^+^, Sca1^+^ CD90^-^, and Sca-1 CD90^+^) for D2.0R cell line. (**d**) Quantification of sphere formation for each population (ALDH^-^ CD90-Sca-1-, ALDH+, ALDH-CD90-Sca-1+, and ALDH-CD90+) for D2.0R cell line. (**e**) Quantification of sphere formation for non-LRC vs LRC after FACS. P-values were calculated using t test (for normally distributed data) or Wilcoxon test with Bonferroni correction for multiple comparisons when appropriate (* p < 0.05, ** p < 0.01, *** p < 0.001, **** p<0.0001).

In human breast cancer cell lines, the proliferative CSC is more epithelial while the quiescent CSC is more mesenchymal (*13*). We compared the percentage of cells expressing E-cadherin for each CSC marker relative to the total live population of cells (**Fig. 2c**). E-cadherin expression was increased in the ALDH^+^ and Sca-1^+^CD90^-^ cells and decreased in Sca-1^-^CD90^+^ relative to total live cells, suggesting these populations are epithelial and mesenchymal, respectively. We next compared the sphere formation efficiency and proliferation of each population after FACS sorting. Sphere forming efficiency was higher for all populations relative to the ALDH^-^CD90^-^Sca-1^-^ cells (**Fig. 2c**), that CSC properties are enriched across all CSC populations tested. Additionally, we found that LRC also demonstrated increased sphere formation relative to non-LRC (**Fig. 2d**). We also monitored the proliferation of each sorted population in normal growth media conditions (10% FBS) by measuring the percent confluence overtime with the Incucyte imaging system (**Fig. S2**). Consistent with other results, ALDH^+^ cells had the fastest proliferation while Sca-1^-^CD90^+^ cells had the slowest proliferation. Finally, we performed limiting dilution, tumor initiation assays for ALDH^+^/^-^ and bulk cells as the gold standard for CSC activity. We found the TIC frequency was highly dependent on the mouse strain used, though ALDH^+^ had the highest TIC frequency in NODscid and NSG mice (**Fig. S4**). We were unable to perform these experiments with CD90^+^ cells as we found the antibodies labeling the cells enhanced NK cell killing of tumor cells via antibody dependent cellular cytotoxicity both *in vitro* and *in vivo*.

### Immune dependence of long-term dormancy

After determining that PyMT, Met-1, and D2.0R cell lines display metastatic dormancy *in vivo* after intracardiac inoculation, we next investigated the importance of the immune system in this phenotype. We performed intracardiac inoculation of each cell line in hosts differing in immunocompetence: fully immunocompetent (FVB/N for PyMT and Met-1, Balb/c for D2.0R), as well as immunodeficient NODscid, or NSG mice (**Fig. 3a**). NODscid mice lack T and B cells but have NK cells and macrophages, which may be slightly less functional than in the syngeneic host (*14*). NSG mice lack T, B and NK cells. Relative to NODscid mice the primary alteration is the loss of NK cells because of the added mutation of the IL-2 receptor gamma chain (*15*). For PyMT, we found a significant decrease in survival between FVB/N and NODscid mice, and NODscid compared to NSG (**Fig. 3b**). This suggests that metastatic dormancy in PyMT is controlled partly by the adaptive immune system, and partly by an IL-2 receptor dependent population, likely NK cells. For Met-1, we found a significant decrease in survival between FVB/N and both NODscid and NSG mice, but no difference between NSG and NODscid (**Fig. 3c**), suggesting that only the adaptive immune system plays a role in Met-1 dormancy. For D2.0R, we found a significant decrease in survival between Balb/c and NSG, but no significant difference between NODscid and Balb/c (**Fig. 3d**), suggesting that the adaptive immune system is not required for metastatic dormancy of D2.0R, but rather a population of immune cells that is IL-2 receptor dependent controls dormancy. Taken together, these findings indicate that the immune system plays a role in metastatic dormancy in all three syngenetic mouse models, but the specific immune cell populations that control dormancy differ between models.

**Figure 3:**
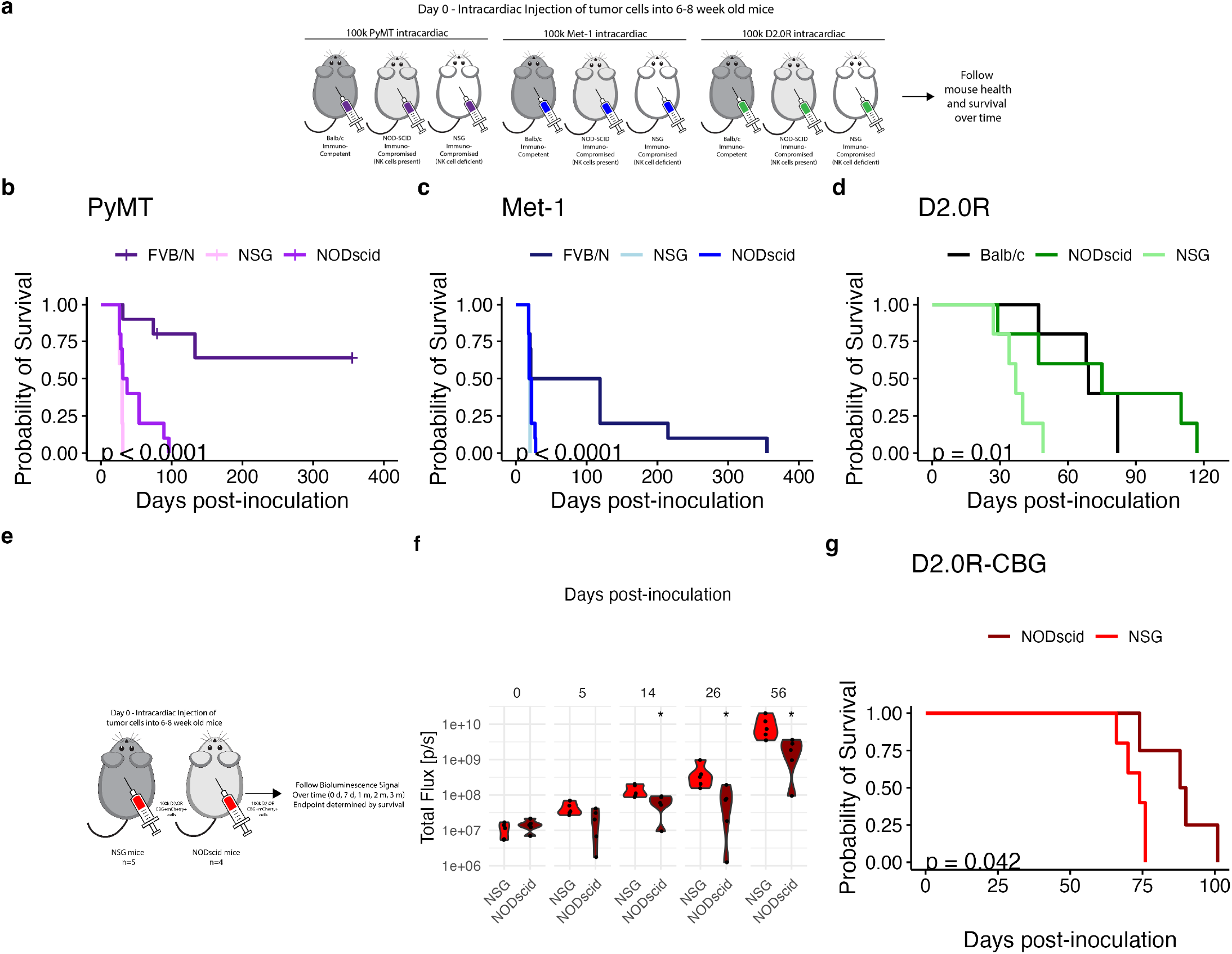
PyMT, Met-1, and D2.0R metastatic dormancy is dependent upon the immune system. (**a**) Schematic for long-term dormancy assays *in vivo* to test the role of the immune system in dormancy maintenance by comparing survival times for syngeneic mice versus NSG (no T, B, nor NK cells) and NODscid (no T, B cells) for each cell line. (**b**) Evaluation of survival for mice (FVB/N, NSG or NODscid) injected with 100,000 PyMT cells. (**c**) Evaluation of survival for mice (FVB/N, NSG or NODscid) injected with 100,000 Met-1 cells. (**d**) Evaluation of survival for mice (Balb/c, NSG or NODscid) injected with 100,000 D2.0R cells. (**e**) Schematic for long-term dormancy assay to quantify bioluminescence over time and survival for mice injected with 100,000 D2.0R cells expressing click beetle green luciferase (CBG). (**f**) Bioluminescence quantification (total flux per animal) over time in NODscid or NSG mice. (**g**) Survival of NODscid or NSG mice injected intracardiac with 100,000 D2.0R-CBG+ cells. Survival curves were compared via log-rank test calculation of p-value. P-values were calculated using t test with Bonferroni correction for multiple comparisons when appropriate (* p < 0.05, ** p < 0.01, *** p < 0.001, **** p<0.0001).

We focused on the D2.0R model in which all mice succumbed to metastasis after a period of dormancy. To confirm that survival times differ between NODscid and NSG for mice injected with D2.0R cells we used click-beetle green luciferase (CBG) D2.0R cells. We injected 100,000 luciferase-expressing D2.0R tumor cells into the left ventricle of NODscid or NSG mice and measured bioluminescence over time using IVIS (**Fig. 3e**).

The bioluminescent signal was initially the same but diverged between NODscid and NSG starting at 14 days and maintained separation through 56 days following injection (**Fig. 3f**, **Fig. S5**). This indicates that initial clearing of tumor cells does not change in NODscid compared to NSG, but rather their ability to grow after metastatic seeding is impaired in NODscid mice. We also confirmed that there was a significantly shorter survival time for NSG mice relative to NODscid for mice injected with D2.0R-CBG (**Fig. 3g**). These results confirm that an IL-2 receptor dependent population of immune cells controls metastatic dormancy of D2.0R cells.

To determine which immune cell population was responsible for these differences in dormancy we determined the effect of depletion of specific immune cell populations. We hypothesized that NK cells were the IL-2 receptor dependent population since they are present in NODscid but absent in NSG mice. However, others have reported that macrophage function may also be altered by the loss of IL-2 receptor in NSG mice (*16*). We tested the importance of both NK cells and macrophages via depletion with Anti-ASGM1 (**Fig. 4a**) or Clodrosome ® (**Fig. 4d**) respectively. Our depletion strategy was successful in both cases (**Fig. S6**), and we measured tumor growth by bioluminescent signal over time in depleted and control mice. Signal was significantly increased in mice depleted of NK cells (**Fig. 4b**), and their survival time was significantly shorter (**Fig. 4c**). However, for mice with macrophage depletion (Clodrosome ®), there was no significant difference in bioluminescent signal over time relative to control (**Fig. 4e**), nor in survival times (**Fig. 4f**). These results indicate that NK cells are primarily responsible for the metastatic dormancy phenotype in the D2.0R model.

**Figure 4:**
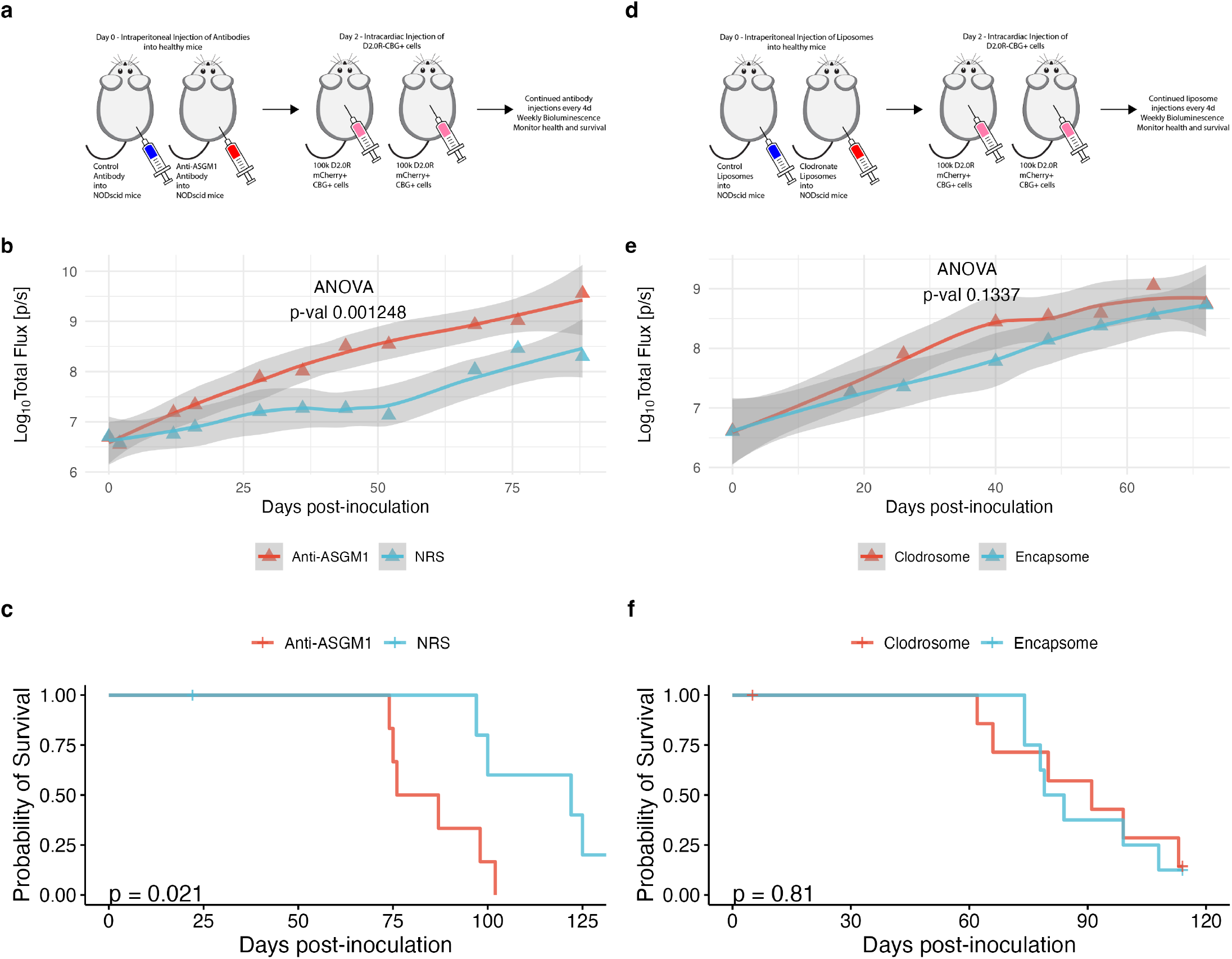
D2.0R metastatic dormancy is partially controlled by NK cells. (**a**) Schematic for long-term dormancy assay in vivo to test the role of NK cells in dormancy maintenance by comparing survival times for mice injected with 100,000 D2.0R-CBG+ cells intracardiac and treated with Anti-Asialo GM1 (Anti-ASGM1) to deplete NK cells or normal rabbit serum (NRS) as a control. (**b**) Bioluminescence quantification (log10 of total flux in photons/second) over time in mice inoculated with luciferase-expressing D2.0R cells and treated with NRS or Anti-ASGM1. (**c**) Evaluation of survival times for mice inoculated with 100,00 D2.0R-CBG+ cells intracardiac and treated with Anti-ASGM1 or NRS. (**d**) Schematic for long-term dormancy assay in vivo to test the role of macrophages in dormancy maintenance by comparing survival times for mice injected with 100,000 D2.0R-CBG+ cells intracardiac and treated with clodrosome to deplete macrophages or encapsome as a control. (**e**) Bioluminescence quantification (log10 of total flux in photons/second) over time in mice inoculated with luciferase-expressing D2.0R cells and treated with encapsome or clodrosome. (**f**) Evaluation of survival times for mice inoculated with 100,00 D2.0R-CBG+ cells intracardiac and treated with encapsome or clodrosome. P-values were calculated using ANOVA. Survival curves were compared via log-rank test calculation of p-value.

### Proliferative CSCs are sensitive to natural killer cell cytotoxicity while quiescent CSCs are resistant

We next conducted NK cytotoxicity assays to examine the sensitivity of CSC populations to NK cells *in vitro*. We first validated that all cell lines are sensitive to NK cell killing via comparing cytotoxicity among all cell lines to the model NK-sensitive cell line Yac-1 (**Fig. S7a**). We also tested the impact of NK cell co-culture on colony formation and found D2A1, D2.0R, and PyMT cell lines had reduced colony formation after treatment with NK cells, but Met-1 did not (**Fig. S7b**). Next, we performed indirect cytotoxicity assays (**Fig. 5a**), where we co-cultured D2.0R cells and NK cells and compared the populations relative to monoculture. In the direct (**Fig. 5e**) assays, we sorted specific CSC populations via FACS and then co-cultured them with NK cells. We first stained untreated (target only) or NK treated (target + NK) for CSC markers (**Fig. 5b**) Sca-1^+^CD90^-^ cells were significantly decreased, suggesting that this population is sensitive to NK cells. We also stained for EMT markers and found EpCAM expressing cells decreased while Vimentin expressing cells increased upon treatment with NK cells (**Fig. 5c**). Finally, LRC percentage was significantly enriched when D2.0R cells were cultured with NK cells (**Fig. 5d**, see **Fig. S8** for gating scheme). Taken together the indirect assays suggest that Sca-1^+^CD90^-^, epithelial, proliferative D2.0R cells are more sensitive to NK cells than their Sca-1^-^CD90^+^, mesenchymal, quiescent counterparts.

**Figure 5:**
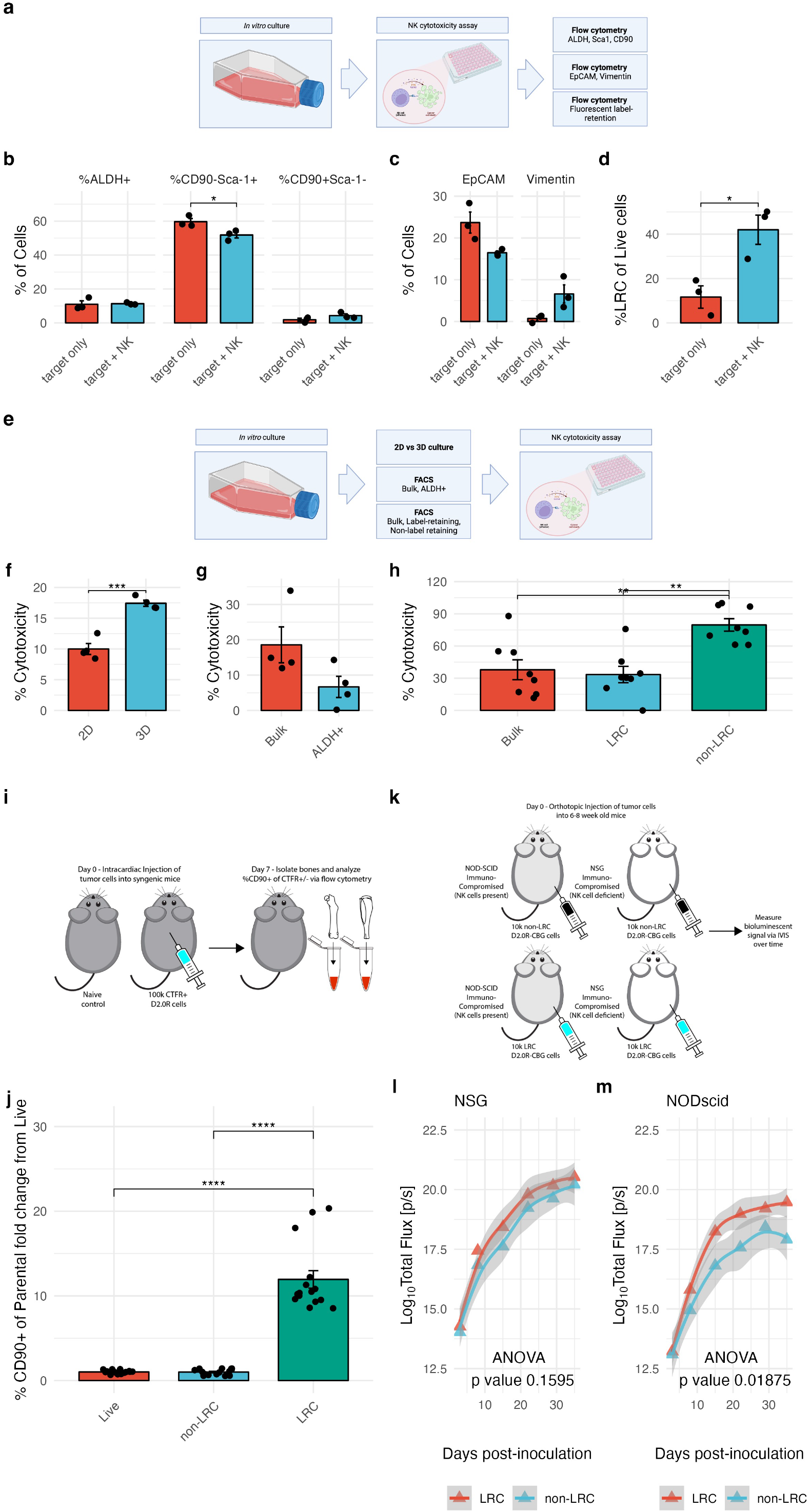
Quiescent D2.0R cells are more resistant to NK cytotoxicity compared to proliferative D2.0R cells *in vitro* and *in vivo*. (**a**) Experimental schematic for indirect analysis of NK cytotoxicity via performing NK co-culture cytotoxicity assay and evaluating cancer stem cell markers, EMT markers, and fluorescent label-retention with and without NK cells via flow cytometry. (**b**) Flow cytometry evaluation of ALDH+, CD90+Sca1-, and CD90-Sca1+ as a percentage of live cells in tumor cells cultured alone (target only) or cultured with NK cells (target + NK). (**c**) Flow cytometry evaluation of EpCAM and Vimentin as a percentage of live tumor cells cultured alone (target only) or co-cultured with NK cells (target + NK). (**d**) Flow cytometry evaluation of LRC as a percentage of live tumor cells cultured alone (target only) or co-cultured with NK cells (target + NK). (**e**) Experimental schematic for direct analysis of NK cytotoxicity via performing 2D or 3D culture, fluorescence activated cell sorting (FACS) for ALDH+, and FACS for LRC D2.0R cells before NK cytotoxicity assay analysis via CytoTox96 quantification of LDH. (**f**) Percent NK-driven cytotoxicity of D2.0R cells cultured in 2D (2% FBS and tissue culture treated flasks) and 3D (mammosphere media and ultra-low attachment flasks) measured by LDH quantification with CytoTox96. (**g**) Percent NK-driven cytotoxicity of FACS sorted Bulk and ALDH+ D2.0R cells measured by LDH quantification with CytoTox96. (**h**) Percent NK-driven cytotoxicity of FACS sorted Bulk, LRC, and non-LRC D2.0R cells measured by LDH quantification with CytoTox96. (**i**) Experimental schematic for analysis of CD90+ expression and maintenance of label retaining D2.0R cells after intracardiac inoculation. (**j**) Flow cytometry quantification of CD90 marker-expression as a percentage of parental (Live, non-LRC, or LRC) D2.0R cells isolated from bone marrow. (**k**) Experimental schematic for analysis of sensitivity of non-LRC) and LRC sensitivity to NK cells and growth in vivo after inoculation of 10,000 non-LRC or LRC D2.0R-CBG+ cells into the fourth left and right mammary fat pads of NSG or NODscid mice and monitored over time with bioluminescence. (**i**) Quantification of bioluminescent signal over time for NSG mice inoculated with 10,000 non-LRC or LRC D2.0R-CBG+ cells. (**m**) Quantification of bioluminescent signal over time for NODscid mice inoculated with 10,000 non-LRC or LRC D2.0R-CBG+ cells. P-values were calculated using t test (for normally distributed data) or Wilcoxon test with Bonferroni correction for multiple comparisons when appropriate (* p < 0.05, ** p < 0.01, *** p < 0.001, **** p<0.0001). Schematic figures made with Biorender (panels a, e).

To confirm these results, we performed assays directly measuring the sensitivity of D2.0R cells in 2D or sphere culture (3D). We previously identified that low serum 2D culture enriches Sca-1^-^CD90^+^ cells relative to sphere culture, due to the low serum conditions enriching for quiescent cells (**Fig. 2a**). We found that cells cultured in 2D were less sensitive to NK lysis than those cultured as spheres (**Fig. 5f**). We next sorted ALDH^+^ or bulk (mixed population) D2.0R cells from 2D culture conditions and found no difference in cytotoxicity (**Fig. 5g**). Finally, we sorted bulk (mixed population), LRC, and non-LRC and found that non-LRC were significantly more sensitive to NK lysis (**Fig. 5h**). In summary, these results support the indirect assays and suggest that Sca-1^+^CD90^-^, epithelial, proliferative non-LRC D2.0R cells are more sensitive to NK cells than their Sca-1^-^CD90^+^, mesenchymal, quiescent LRC counterparts.

We next determined whether the findings from our *in vitro* experiments also apply in the *in vivo* setting. We injected CTFR-labeled D2.0R cells into the left ventricle of balb/c mice and isolated bone marrow after 7 days (**Fig. 5i**). Using CD90 staining, we analyzed the overlap between CD90 expression and label retention *in vivo*. We compared the CD90 expression in live tumor cells to non-LRC and LRC and found a significant increase in CD90 expression in the LRC population relative to both live and non-LRC (**Fig. 5j**). This result supports our *in vitro* findings, indicating that the CD90^+^ population is predominately label retaining *in vivo*. We also investigated the resistance of LRC D2.0R cells to NK cells *in vivo*. We sorted non-LRC and LRC using FACS and inoculated them into the mammary fat pads of either NODscid or NSG mice and monitored tumor growth using IVIS (**Fig. 5k**). We found no significant difference in the growth rate of non-LRC versus LRC in NSG mice (**Fig. 5l**). However, in NODscid mice, which have NK cells, LRCs demonstrate a significant growth advantage relative to non-LRC (**Fig. 5m**). This finding suggests that LRC are more resistant to NK cell-mediated lysis *in vivo* compared to non-LRC.

### BACH1 and SOX2 control tumor sensitivity to NK cells

To identify mechanisms linking dormancy to NK cytotoxicity in a non-biased manner, we performed RNA-seq of LRC and non-LRC. 4199 genes were differentially expressed (adjusted p-value < 0.05) between the two cell populations (**Fig. 6a**), 279 of which had an absolute log2 fold change > 2. Upon hierarchical clustering, data clustered by condition and not by batch (**Fig. 6b**). GSEA demonstrated pathways related to inflammation including Neutrophil extracellular trap formation, cytokine-cytokine receptor interaction, complement and coagulation cascades, allograft rejection, and others were enriched in LRC population while in non-LRC, enriched pathways were related to cell cycle including cell cycle and DNA replication among others (**Fig. S9**). Activating NK ligands were more highly expressed in non-LRCs (**Fig. 6c**) while MHC genes were decreased in non-LRCs (**Fig. 6d**). Interestingly, STING and STING targets expression were increased in non-LRC (**Fig. 6e**). These results are consistent with the phenotype we observed both *in vitro* and *in vivo* that non-LRC are more sensitive to NK cell cytotoxicity than LRC.

**Figure 6:**
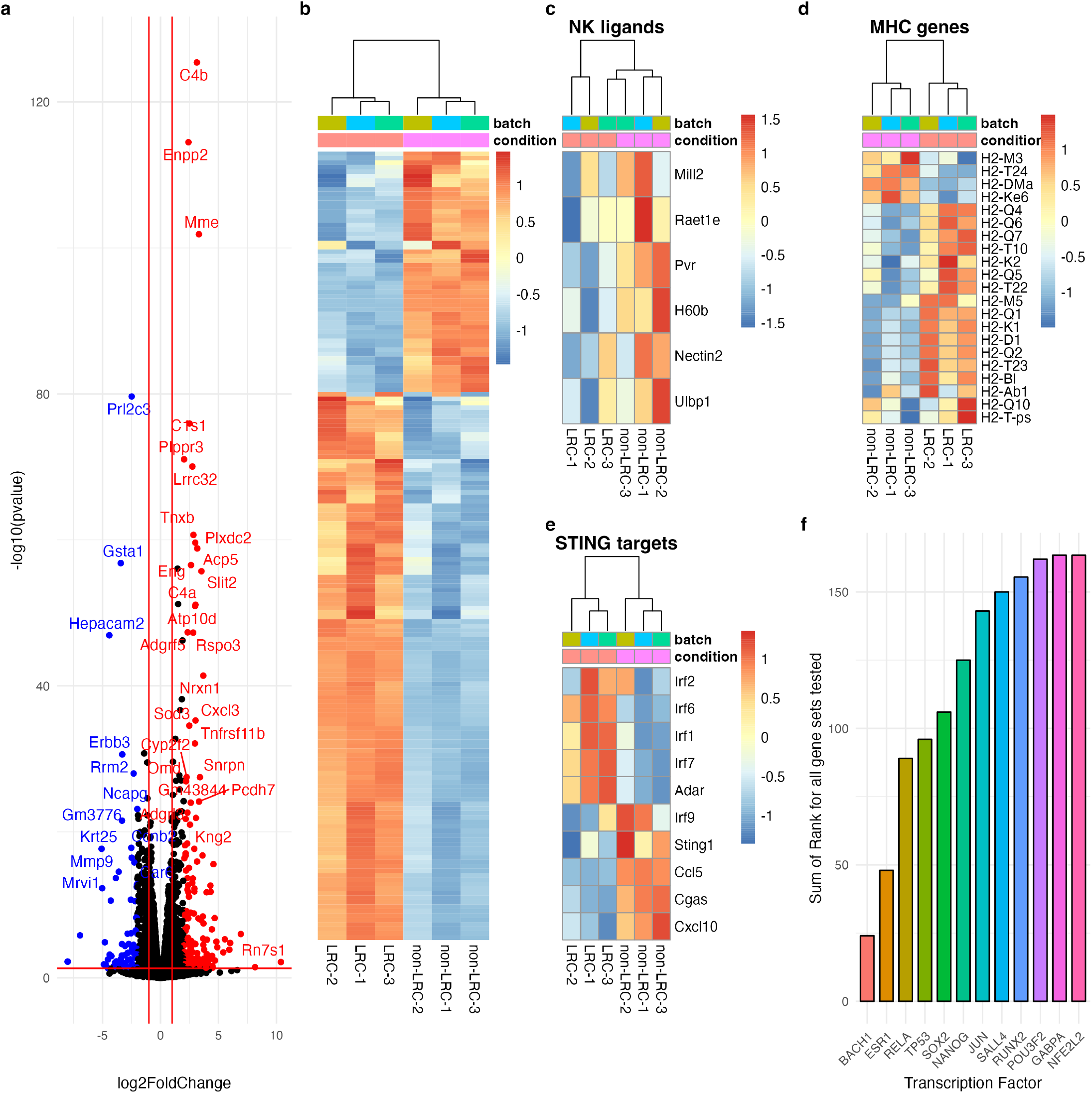
Bulk RNA sequencing of LRC and non-LRC D2.0R cells identifies drivers of NK cell resistance in LRC. (**a**) Volcano plot of differentially expressed genes up in LRC (red) and non-LRC (blue). (**b**) Heatmap of differentially expressed genes with log2 foldchange greater than 2 and adjusted p value < 0.05. (**c**) Heatmap of NK ligand gene expression. (**d**) Heatmap of MHC gene expression. (**e**) Heatpmap of STING target gene expression. (**f**) ChEA3 analysis of predicted transcription factors driving DEGs in LRC, stemness, EMT, complement, and MHC gene expression via sum of ranks of predicted transcription factors for all gene sets tested.

We next investigated transcription factor drivers of these gene expression programs using ChEA3 analysis (*17*). We compared the predicted transcription factors for the genes that were up in LRC to the predicted transcription factors for MHC genes, complement genes, and genes associated with EMT and murine mammary stem cells. We reasoned that the overlap of these gene sets should identify transcription factors that are responsible for general differences in gene expression, as well as for the increase in MHC gene expression and EMT/stemlike phenotype. *Bach1* was the top ranked transcription factor within the intersection of these gene sets (**Fig. 6f**).

Additionally, *Sox2* was the fifth ranked transcription factor and as *Bach1* has been shown to regulate *Sox2* via increasing its stability (*18*) we chose to investigate these genes further.

To confirm these transcription factors as drivers of NK resistance and sensitivity respectively, we knocked these genes down using a Tet-On inducible shRNA system in D2.0R-CBG cells. We found knockdown of Bach1 or Sox2 did not alter proliferation or cell eccentricity (**Fig. S10**). We confirmed that knockdown of Bach1 reduces Sox2 protein levels (**Fig. 7a-b**), as reported by Wei et al (*18*). We also tested whether knockdown of Bach1 or Sox2 altered the sensitivity of D2.0R cells to NK cell cytotoxicity. We found a small but statistically significant increase in cytotoxicity after knockdown of Bach1 and a trend toward increased NK cytotoxicity after knockdown of Sox2 (**Fig. 7c**). Finally, we inoculated these cells into the mammary fat pads of NODscid or NSG mice and treated mice with either control water or doxycycline containing water to investigate the difference is tumor growth. We found knockdown of Bach1 and Sox2 only decreased tumor growth in NODscid mice with NK cells, and not in NSG mice without NK cells (**Fig. 7d-i**). This finding confirms that Bach1 and Sox2 expression contribute to the NK resistant phenotype of D2.0R cells.

**Figure 7:**
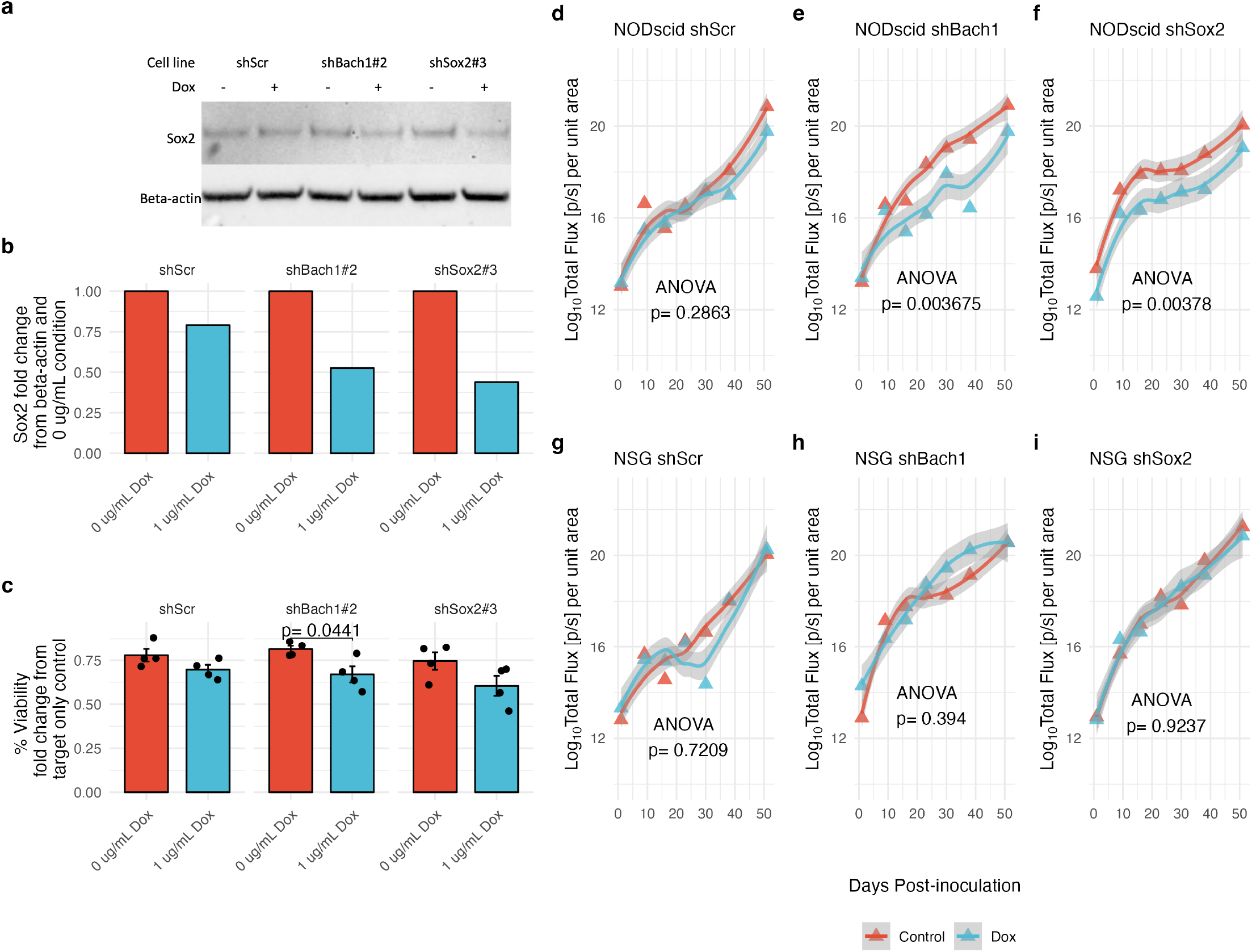
Bach1 and Sox2 expression partially control NK resistant phenotype of quiescent D2.0R cells. (**a**) Western blot for Sox2 and Beta-Actin protein in D2.0R-CBG cells with doxycycline inducible knockdown of shScr (control), shBach1, or shSox2 with or without doxycycline (Dox) treatment. (**b**) Quantification of western blot normalized to beta-actin and 0 ug/mL Dox condition for each cell line. (**c**) NK cytotoxicity for inducible knockdown cell lines D2.0R-CBG shScr (control), shBach1, or shSox2 treated with 0 ug/mL doxycycline or 1 ug/mL doxycycline. (**d-i**) Tumor growth as measured by bioluminescence of D2.0R-CBG cell lines with inducible knockdown of shScr (control), shBach1, or shSox2, in mice that have NK cells (NODscid, d-f) or do not have NK cells (NSG, g-i) comparing tumor growth over time via ANOVA between mice treated with control water or water containing doxycycline over the course of the experiment. P-values were calculated using t-test (normally distributed data) or Wilcox test with Bonferroni correction when appropriate. For tumor growth over time, p-values were calculated using ANOVA.

**Figure 8:**
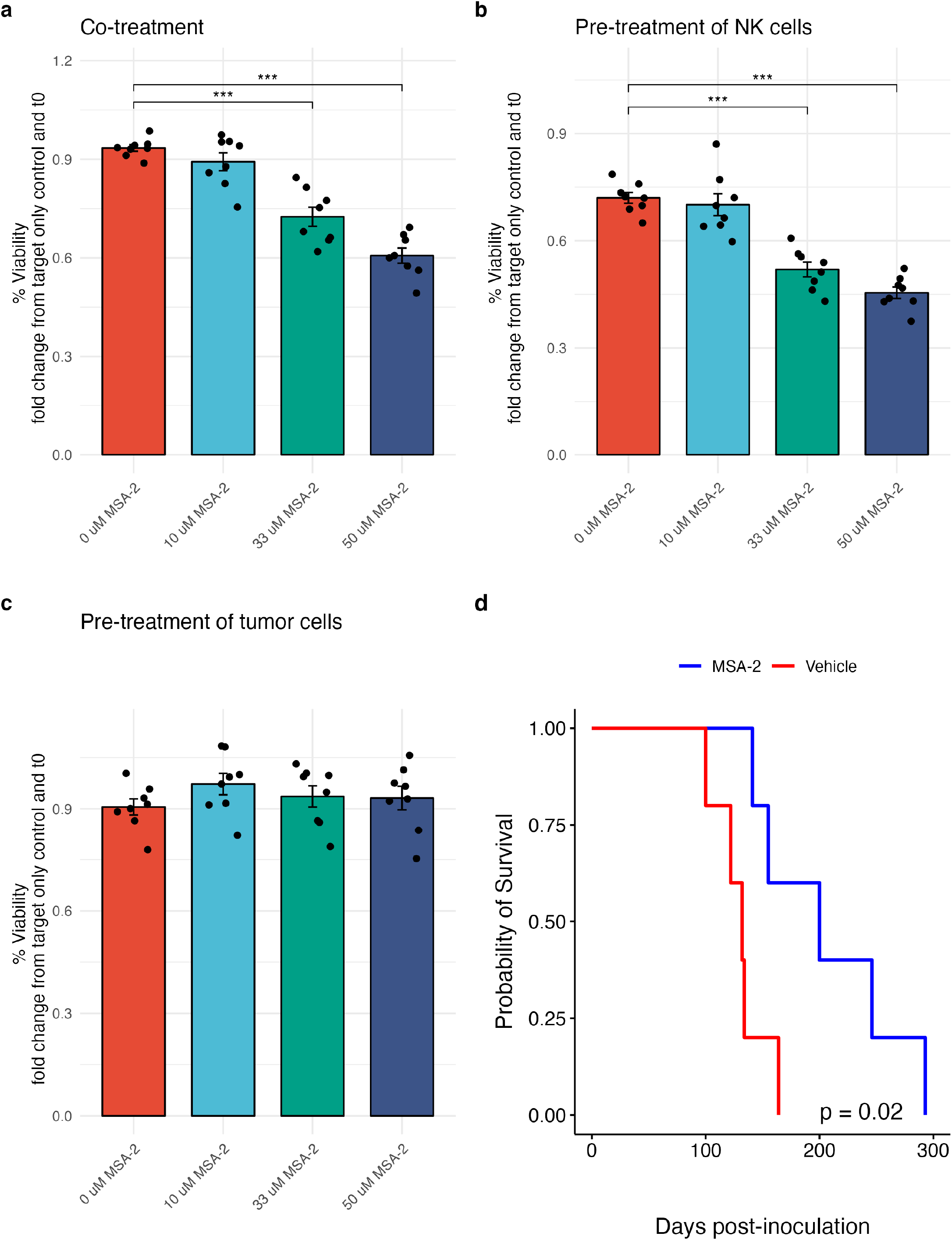
STING agonist MSA-2 increases NK cell killing in vitro and increases survival of mice inoculated with D2.0R cells intracardiac *in vivo*. (**a**) MSA-2 co-treatment increases NK cell killing in vitro with D2.0R cells (**b**) MSA-2 pre-treatment of NK cells increases killing of D2.0R cells in vitro (**c**) Pre-treatment of tumor cells with MSA-2 does not alter NK cell killing (**d**) MSA-2 increases survival in vivo after one treatment with 1 mg MSA-2 7 days after intracardiac inoculation of 100,000 D2.0R tumor cells into balb/c mice. P-values were calculated using t test with Bonferroni correction for multiple comparisons when appropriate (* p < 0.05, ** p < 0.01, *** p < 0.001, **** p<0.0001). For survival Kaplan-Meier curves, p-values were calculated using log-rank test.

### STING agonist MSA-2 increases NK cell killing

Finally, as we found low STING and STING target expression in LRC compared to non-LRC, we investigated the STING agonist MSA-2 as a method to reverse the NK resistant phenotype. MSA-2 was recently reported by the Massagué group to prevent reactivation from dormancy (*19*). MSA-2 significantly enhanced NK cell killing *in vitro* upon co-treatment of tumor cells and NK cells (**Fig. 7a**), or by pre-treatment of NK cells alone (**Fig. 7b**), but not by pre-treatment of tumor cells alone (**Fig. 7c**). This indicates in this system STING activation in NK cells but not in tumor cells mediates the enhanced killing observed. This is contrary to the Kpad1 model reported by the Massagué lab (*19*) in which tumor STING activation predominated. We also found that a single dose of MSA-2 given 7 days after inoculation significantly increased mouse survival time. (**Fig. 7d**).

## Discussion

Understanding breast cancer dormancy has emerged as a major research focus in the last decade (*20*). While much of this research has focused on identifying tumor-intrinsic mechanisms of breast cancer dormancy, some studies have explored the role of the immune system. The lack of immunocompetent models poses a major challenge in this research effort. Human tumor cells inoculated into immunocompromised hosts have been used in many studies but the species mismatch between the host and the tumor cells is a major limitation. Although interactions with innate immune cells are still present in this model and may be crucial for the dormancy phenotype, it does not reflect the clinical setting. In this study, we evaluated the metastatic dormancy phenotype of five murine mammary carcinoma cell lines and identified three lines that exhibited long-term metastatic dormancy, with survival exceeding one hundred days following the intracardiac inoculation of 100,000 tumor cells.

Cancer stem cells (CSCs) have been extensively studied in human breast cancer cell lines since their initial identification by Al Hajj et al. in 2003 (*21*). However, CSCs have been less well studied in murine models. One reason for this is that the CSC hypothesis was called into question after a report from Sean Morrison’s group demonstrated that, in melanoma, the tumor initiation ability of a cell population was highly dependent on the immune background used (*22*). Although only a subpopulation of melanoma cells was capable of tumor initiation in NOD/SCID mice, NSG mice without NK cells were able to form tumors from single melanoma cells, calling into question the relevance of the cancer stem cell hypothesis. However, our present study as well as others have now shown that immune evasion is an important characteristic of CSC’s and stress the importance of using immunocompetent models to generate clinically relevant data. Two CSC populations have been reported in PyMT, a tumor-initiating (ALDH^+^) and a metastasis-initiating (CD24^+^CD90^+^) population (*23*). We chose CD90 and ALDH as markers to investigate for all cell lines because of this report. We also chose Sca-1 as it was reported to be a CSC marker in TUBO, PyMT, 4T1, and other murine cell lines (*10*). Our findings indicate that the ALDH^+^ and Sca-1^+^CD90^-^ populations were more proliferative and epithelial, while the Sca-1^-^CD90^+^ population was more quiescent and mesenchymal, in agreement with the report in PyMT (*23*) and the dichotomy seen in human breast cancer cells (*13*).

NK cells play a crucial role in determining the outcome of metastasis (*24*). The earliest studies on metastasis by Hanna et al in 1981 showed that NK cells cleared tumor cells after intracardiac inoculation (*4*). This classic study has recently been revisited using modern techniques such as bioluminescence and intravital imaging (*25*). The importance of NK cells in metastatic dormancy has also been established. Malladi et al. found that human dormancy-competent cell lines required NK cells for the dormancy phenotype (*5*). However, one disadvantage of using human tumor cells in mouse models is that there is a species mismatch between human tumor cells and mouse NK cells. This mismatch can alter the strength of activating and inhibitory receptor interactions and may result in reduced activation or inhibition (*26*). The interactions between dormancy-competent cell lines and NK cells were validated using human NK cells and tumor cells *in vitro* (*5*), but this approach does not replicate the *in vivo* metastasis environment. In this report, we used syngeneic interactions between D2.0R tumor cells and NK cells *in vitro* and *in vivo* to validate the importance of NK cells in D2.0R metastatic dormancy.

We found that the quiescent, mesenchymal D2.0R CSC population is resistant to NK lysis, while the epithelial, proliferative D2.0R CSC population is sensitive to NK lysis. This is the first report to investigate the sensitivity of D2.0R cells to NK cells and is consistent with previous literature that showed that TUBO spheres are sensitive to NK lysis while cells grown in 2D are resistant (*11*). In contrast, there are multiple reports that studied the sensitivity of human breast cancer CSC to NK lysis. Human BC has a quiescent, mesenchymal CSC population (CD24^-^CD44^+^) and a proliferative, epithelial CSC population (ALDH^+^) (*13*). Three reports tested the sensitivity of the CD24^-^CD44^+^ population to NK cells and two of three found they were more resistant to NK cells than their differentiated counterparts (*27–29*). Similarly, three reports tested the sensitivity of the ALDH^+^ population to NK cells and two of three found they were more sensitive to NK cells than their differentiated counterparts (*11, 30, 31*). Moreover, co-culture of NK cells and breast cancer cells increases the mesenchymal stem-like population (*32*). Interestingly, EMT and stemness have been shown to decrease sensitivity of tumor cells to cytotoxic T cell and NK lysis (*33*). This phenomenon may hold across mouse and human breast cancers, but for other cancers, there is some evidence that EMT enhances sensitivity to NK cell killing and stem-like cells are more sensitive to NK lysis than their differentiated counterparts (*34*). Thus, it is not yet clear how widely applicable this model may be.

In this report, we have identified three immunocompetent models of metastatic dormancy with varying immune dependencies. We have conducted a detailed investigation of the D2.0R model and found that metastatic dormancy in this model is dependent on NK cells, with different sensitivity of quiescent mesenchymal CSCs compared to proliferative epithelial CSCs, which is influenced by RAE1 expression. To prevent disease recurrence, it may be possible to target the mechanisms that lead to quiescent CSC resistance to NK cells. In addition, we demonstrate that this quiescent CSC population may be sensitive to STING agonists. The utilization of immunocompetent mouse models may facilitate the development of strategies to effectively target dormant CSC reducing metastatic recurrence, a major cause of breast cancer mortality.

## Materials and Methods

### Cell culture

#### Murine mammary carcinoma cell lines

Met-1 cells were a gift from the laboratory of Alexander D Borowsky (*35*). D2.0R and D2A1 cells were derived by the Jeffrey Green lab and were a gift from the laboratory of Hunter Kent (*36*). PyMT cells were derived from a spontaneous MMTV-PyMT tumor in the laboratory of Max S Wicha. E0771 cells were a gift from the laboratory of Qiao Li.

#### Maintenance of cell lines

Murine mammary carcinoma cell lines were cultured at 37’C and 5% CO2 in a humidified incubator. PyMT and Met-1 cells were maintained in RPMI 1640 (Gibco) with 10% FBS. D2.0R, D2A1 and E0771 were maintained in DMEM (Gibco) with 10% FBS and 1x GlutaMAX (Gibco). Cells were passed with TryplE (Gibco) and subcultured at a 1:20 ratio upon reaching 90% confluency. Cells were confirmed mycoplasma-free every two months via MycoAlert Mycoplasma Detection Kit (Lonza).

#### 2D and sphere culture

Murine mammary carcinoma cell phenotypes were tested in 2D and sphere culture conditions. 2D culture was defined as attachment culture with low serum 2% FBS that supported slow growth over a period of one week. sphere culture was defined as ultra-low attachment culture with mammosphere media that supported formation of tumor spheres over a period of one week. Mammosphere media was prepared with 475 mL of MEBM (Lonza) with 1x GlutaMAX (Gibco), 1x B27 (Gibco), 1x Pen/Strep (Gibco), 5 ug/mL insulin (Sigma), 20 ng/mL EGF (BD Biosciences), bFGF (Gibco), 0.5 ug/mL hydrocortisone (Sigma), 0.4 IU/mL heparin (StemCell), and 1% wt/vol of 400 cP methylcellulose (Sigma). Mammosphere media was prepared and placed on a rocker overnight at 4’C to dissolve methylcellulose. After the methylcellulose was fully dissolved, media was sterile filtered and stored at 4’C for up to two months.

#### Fluorescent dye labeling for label retention assays

PKH67 green, fluorescent cell linker midi kit (Sigma) was used according to manufacturer instructions to label murine mammary carcinoma cell lines. CellTrace Far Red (CTFR) Cell Proliferation kit (Thermo Fisher) was used according to manufacturer instructions to label murine mammary carcinoma cell lines. For short-term dormancy and immunogenicity studies *in vivo*, tumor cells were labeled with fluorescent dye and immediately injected into mice. For studies that investigated the differential phenotype of proliferative versus quiescent tumor cells, cells were labeled with fluorescent dye and cultured for one week in 2D conditions described above to generate a label retaining population that was quiescent and a non-label retaining population that was proliferative.

#### Sphere formation assay

For sphere formation assays, cells were sorted, and counts were taken from the sorter as described above. Cells were seeded at one hundred cells per well in mammosphere media as described above and allowed to form spheres for one week. After one week, spheres larger than 70 microns were manually counted via light microscopy and recorded.

#### Sphere formation assay

For colony assay, tumor cells were co-cultured with primary NK cells for 24 hr. Primary NK cells were isolated from spleens of naïve balb/c mice at 6-10 weeks of age. Splenocytes were isolated via mashing against a 70-micron cell strainer and red blood cells were removed via five seconds of hypotonic lysis in water. NK cells were purified from total splenocytes using the NK cell isolation kit, mouse (Miltenyi Biotec) for magnetic activated cell sorting using the MACS system (Miltenyi Biotec). This kit uses a depletion strategy and the negative fraction after sorting is natural killer cells. NK cells were cultured in RPMI 1640 (Gibco) with 10% FBS (Gibco), 0.375% sodium bicarbonate (Gibco), 1 mM sodium pyruvate (Gibco), 1% pen/strep (Gibco), 1000 U/mL murine recombinant IL-2 (PeproTech), and 0.001% 2-mercaptoethanol (Sigma) and used immediately for co-culture assays. Target cells were first passaged from original culture conditions, counted using the Luna slide counter (Logos Biosystems) with acridine orange and propidium iodide identification of live and dead cells respectively and resuspended at 1e6 cells/mL in PBS. Cells were then diluted to 1e5 cells/mL in NK media and hundred microliters of cells were plated with two fifty microliters of NK cells in 6 well plate. After 24 hr, culture media was aspirated, and tumor cells were washed with PBS two times to remove any non-adherent NK cells. Adherent tumor cells were trypsinized, resuspended in tumor cell media and counted by luna slide counter. Briefly, Tumor cells survived by Nk cell cytotoxicity and untreated control tumor cells were were inoculated in 6-well plates (500 cells/well) and incubated for 8 days. The colony formation of cells was determined using crystal violet staining (Beijing Solarbio Science & Technology Co., Ltd. Cells were then fixed with 100% methanol for 10 min at room temperature. Next, cells were stained by 0.1% crystal violet for 5 min at room temperature. Finally, the cell colonies were observed under an inverted microscope (IX73; Olympus Corporation, Tokyo, Japan; magnification, ×2.5) and manually counted.

### Proliferation quantification with Incucyte

For measurement of proliferation over time, cells were sorted and seeded at one hundred cells per well in full serum media and allowed to grow for two weeks in an Incucyte microscope incubator. Images were taken every 6 hours and the confluency mask was used to calculate the percent confluency over time. Data was exported to csv format from Incucyte software and analyzed using R.

#### Animal Studies

All animal studies were approved by the University of Michigan Institutional Animal Care and Use Committee (IACUC) and were performed in accordance with approved protocols.

#### Intracardiac inoculation

Tumor cells were prepared in sterile PBS such that the number of cells desired to be delivered to each mouse was in a volume of one hundred microliters. Mice were anesthetized with isoflurane and the chest wall was depilated and cleaned with alcohol. A bright light was used to localize the heartbeat through the chest wall and the bottom right area of this region was taken to be the left ventricle. An insulin syringe was prepared with a 100-microliter air gap and 150 microliters of cell suspension was drawn into the syringe and bubbles were removed. The needle was slowly inserted into the chest at the site of the heartbeat and correct placement in the left ventricle was confirmed by a bright flash of pulsing blood into the syringe. Following proper placement, one hundred microliters was expelled from the syringe and then slight negative pressure was applied to prevent tumor cells spilling into the chest cavity. Pressure was applied to the chest wall and mice were returned to a clean cage and monitored until they recovered from anesthesia.

#### Survival monitoring

Mice that received tumor cells were monitored at least twice weekly for the duration of the experiment. Mice were euthanized once they reached moribund conditioning (greater than 20% decrease in body weight, limited physical activity, labored breathing, significant hunching behavior, or limping).

#### Orthotopic inoculation

Tumor cells were prepared in sterile PBS with 50% Matrigel (Corning) such that the number of cells desired to be delivered to each injection was in a volume of fifty microliters. Mice were anesthetized with isoflurane and the skin above the fourth mammary fat pads was depilated and cleaned with alcohol. A bright light was used to localize the fourth nipple and the mammary fat pad was grabbed using Adson skin grabbers near the nipple. An insulin syringe was used to deliver fifty microliters of cell suspension to the fat near the nipple through the skin. A successful injection was confirmed by a ballooning of the fat at the injection site. Mice were returned to a clean cage and monitored until they recovered from anesthesia.

#### Isolation of bone marrow

Mice were euthanized with CO_2_ and euthanasia was confirmed via pneumothorax. The femur and tibia were removed together from the hip joint and cleaned of skin and muscle. The metaphysis at the knee was removed for both the bottom of the femur and top of the tibia to expose the bone marrow cavity. Bones were placed marrow side down in a 0.5 milliliter tube with a small hole in the bottom. This tube was placed in a larger 1.5 milliliter tube and centrifuged at 10,000 x g for 15 seconds to remove the bone marrow from the bones. Successful centrifugation was confirmed by a large red pellet at the bottom of the larger tube and a white appearance of the bones in the top tube. Bones were discarded and marrow resuspended in sterile PBS for further staining or directly proceeding to flow cytometry analysis of label retaining cells.

#### *In vivo* monitoring of bioluminescence

For *in vivo* monitoring of bioluminescence, mice were injected intraperitoneally with d-luciferin (Regis Technologies) at a final dose of 150 mg/kg. Mice were then anesthetized using isoflurane and arranged on a warmed imaging stage in the IVIS Lumina imaging system (Perkin Elmer). After 10 minutes, mice were imaged using “Auto” settings with an open emission filter and the excitation filter blocked. Regions of interest (ROIs) were drawn using Aura software (Spectral Instruments Imaging) and the same ROIs were applied to all subsequent time points. Data was exported as flux (photons/sec) for each ROI and further analyzed using R.

#### In vivo immune cell depletion studies

For depletion of natural killer cells, mice were injected intraperitoneally with three hundred micrograms of Ultra-LEAF purified anti-Asialo GM1 rabbit polyclonal antibody (clone Poly21460, BioLegend) every four days for the duration of the experiment. Control mice were injected with three hundred micrograms of normal rabbit serum (Abcam) every four days for the duration of the experiment. For depletion of macrophages, mice were injected intraperitoneally with 2 mg of Clodrosome (Encapsula) while control mice were injected with 2 mg of Encapsome (Encapsula) every four days for the duration of the experiment.

### Immunofluorescence and confocal microscopy

For immunofluorescence studies, cells were cultured on sterile glass coverslips in a six well plate. For label retention studies combined with Ki-67 labeling (Anti-Ki-67 clone D3B5, Cell Signaling Technology), cells were cultured in low serum for one week. For estrogen receptor staining (Anti-Estrogen Receptor alpha clone 6F11, Abcam), cells were cultured until 80% confluent. For staining, coverslips were washed with PBS, fixed with 10% neutral buffered formalin, and permeabilized with 0.5% Triton-X-100 in PBS for 3 minutes. Following permeabilization, coverslips were washed with PBS twice. Primary and no primary control staining solutions were prepared with a 1:200 dilution of antibody in PBS containing 1% bovine serum albumin and a 50-microliter droplet of staining solution was added to a fresh piece of parafilm, then the coverslip was inverted on the parafilm surface to incubate with the antibody solution for 20 minutes in the dark. The coverslips were washed twice with PBS and the process was repeated with secondary antibody (1:200) containing DAPI for nuclear staining. Coverslips were washed twice with PBS and mounted using Fluoromount-G (SouthernBiotech) onto glass slides and imaged on Nikon A1Si confocal microscope managed by the BCRF microscopy core (University of Michigan).

### Flow cytometry analysis

#### Label retention analysis

For analysis of label retaining cells, fixed unstained cells and fixed freshly labeled cells were used as negative and positive controls, respectively. Bone marrow was isolated using previously described method and analyzed on Coulter Cytoflex or Bio-Rad ZE5 managed by the flow cytometry core (University of Michigan) using the 488 laser and 525/35 emission filter for PKH67 and the 640 laser and 670/30 emission filter for CTFR. For all experiments, the presence of label-retaining cells was reported as a fold-change from a naïve control to account for any auto-fluorescent background present in bone marrow.

#### CSC panel

For analysis of cancer stem cell markers, tumor cell lines were cultured in 2D or sphere according to the experiment and plated in 96 well plates in triplicate for each single color control, full minus one (FMO), and mastermix sample including unstained, DAPI only (Thermo Fisher), DEAB (StemCell) as a negative control for ALDEFLUOR, ALDEFLUOR (StemCell) only, PE-isotype control for CD90 (Abcam), PE-CD90 (clone G7, Abcam), APC-isotype control for s (BioLegend), APC-Sca-1 (clone D7, BioLegend), FMO-ALDH, FMO-CD90, FMO-Sca-1, and master mix (DAPI, ALDEFLUOR, CD90, Sca-1). Samples were analyzed on the Bio-Rad ZE5 managed by the flow cytometry core (University of Michigan). FCS files were analyzed using FlowJo (Treestar inc.) and/or R using the FlowCore package. Single color controls were used to create a compensation matrix that was applied to all samples. FMOs were used to set gates for each stain and these gates were applied to the mastermix sample.

#### EMT panel

For analysis of epithelial mesenchymal transition markers, tumor cell lines were cultured in 2D or sphere according to the experiment and plated in triplicate in ninety-six well plates as described above in combination with the CSC markers. All EMT markers were used in the AF647 channel, and the same Sca-1 clone described above was used conjugated to PECy7. The following antibodies were used in addition to those described above: AF647-Ecadherin (clone DECMA-1, BioLegend), AF647-EpCAM (clone G8.8, BioLegend), and AF647-Vimentin (clone OD1D3, BioLegend). Samples were analyzed as described above.

#### Fluorescence activated cell sorting

For fluorescence activated cell sorting, cells were stained as described above and sorted using the MoFlo Astrios sorter. Cells were sorted into pure FBS to maintain viability and counts were recorded from the sorting instrument were used to resuspend the cells for downstream *in vitro* or *in vivo* applications.

#### NK cell cytotoxicity assays

For all NK cytotoxicity assays, primary NK cells were isolated from spleens of naïve balb/c mice at 6-10 weeks of age. Splenocytes were isolated via mashing against a 70-micron cell strainer and red blood cells were removed via five seconds of hypotonic lysis in water. NK cells were purified from total splenocytes using the NK cell isolation kit, mouse (Miltenyi Biotec) for magnetic activated cell sorting using the MACS system (Miltenyi Biotec). This kit uses a depletion strategy and the negative fraction after sorting is natural killer cells. NK cells were cultured in RPMI 1640 (Gibco) with 10% FBS (Gibco), 0.375% sodium bicarbonate (Gibco), 1 mM sodium pyruvate (Gibco), 1% pen/strep (Gibco), 1000 U/mL murine recombinant IL-2 (PeproTech), and 0.001% 2-mercaptoethanol (Sigma) and either used immediately for cytotoxicity assays or cultured overnight before adding target cells. Target cells were first passaged from original culture conditions, counted using the Luna slide counter (Logos Biosystems) with acridine orange and propidium iodide identification of live and dead cells respectively and resuspended at 1e6 cells/mL in PBS. If required, cells were sorted via FACS as previously described. Cells were then diluted to 1e5 cells/mL in NK media and fifty microliters of cells were plated with fifty microliters of NK cells in each well of a ninety-six well plate. Target and effector cells were cultured 4 hours or overnight and cytotoxicity was measured using the CytoTox96 LDH assay (Promega) or using click beetle green luciferase expressing cells and read on a BioTek Synergy H1 plate reader (BioTek Instruments). The percent cytotoxicity was calculated via manufacturer instructions as (experimental - effector spontaneous - target spontaneous)/(target maximum - target spontaneous) x 100 for LDH assay, or for CBG expressing cells, by using the fold change in luminescence from the tumor cell only control.

#### siRNA knockdown cells

For NK cytotoxicity assays with siRNA knockdown, target cells were plated at 4e3 cells/well in ninety-six well plates with full serum media overnight. Then siRNA lipid complexes were prepared using Opti-MEM media (Gibco), Lipofectamine RNAiMAX (Invitrogen), and Silencer siRNAs (Negative Control No. 1 or pan-*Raet1* siRNA, Thermo Fisher) and complexes were added to cells according to manufacturer instructions. Cells were cultured for 1-3 days and analyzed for knockdown efficiency via flow cytometry or NK cytotoxicity assay.

#### Antibody treatment

For NK cytotoxicity assays with antibody treatment, target tumor cells and effector NK cells were prepared as described above and one microgram of Anti-NKG2D (clone C7, BioLegend) or Armenian Hamster IgG isotype control (clone HTK888, BioLegend) was added per well. Incubation and LDH assay were performed as described above.

### RNA sequencing

#### Bulk RNA-seq

RNA was isolated using the Qiagen RNeasy Micro kit according to the manufacturer instructions. Concentration was determined via nanodrop and A260/A280 evaluated for quality control before further QC, library prep and sequencing by the Advanced Genomics Core (University of Michigan).

### Data analysis and statistics

Statistical analysis and hypothesis testing was performed in R. All reported values represent the mean and standard error of the mean from 3-9 biological replicates, with at least technical triplicates within each experiment. Whenever possible, all data points are included in graphical representations of data. The method used for hypothesis testing is included in each figure caption.

## Acknowledgements

We acknowledge support from the National Institutes of Health NIH R35 CA197585 (to Max Wicha), Breast Cancer Research Foundation BCRF-18-173 (to Max Wicha), and NIH K99 CA267261 (to Grace Bushnell).

## Data Availability Statement

All data required to evaluate the conclusions of this manuscript are present in the main text and supplemental data. Additional data will be available from the authors upon request.

## Author Contributions Statement

GGB and MSW conceived the project. GGB performed all biological experiments with help from DS, HCW, MZ, TDF, CMH, and SO. GGB analyzed the data. GGB wrote the first draft of the manuscript and all authors provided feedback on the manuscript. GGB and MSW critically edited the final draft of the manuscript. GGB and MSW provided funding for the experiments in the manuscript.

