## Supplemental Figures for "Natural killer cell regulation of breast cancer stem cells mediates metastatic dormancy"

**Figure S1: Fluorescent and luminescent protein expression is not compatible with dormancy models, but fluorescent cell membrane/cytoplasm dyes are non-immunogenic. (a)** Comparison of tumor growth for PyMT or PyMT-RFP cell lines in FVB/N immunocompetent or NSG immunocompromised mice. **(b)** Comparison of tumor growth for unlabeled, PKH, or CTFR labeled PyMT cells and PyMT-RFP cells in fully immunocompetent FVB/N mice. **(c)** Comparison of tumor growth for unlabeled or PKH labeled D2.0R cells to D2.0R-Luc-mCherry cells in fully immunocompetent Balb/c mice.

**Figure S2: Murine cell lines contain an epithelial, proliferative cancer stem cell and a mesenchymal, quiescent cancer stem cell population. (a)** Flow cytometry evaluation of cancer stem cell markers ALDH (top), Sca-1$^{-}$CD90$^{+}$ (middle) and Sca-1$^{+}$CD90- (bottom) for PyMT and Met-1 cell lines in 2D (2% FBS tissue-culture treated flasks) and 3D (mammosphere media ultra-low attachment flasks) reported as fold change from 2D. (**b)** Flow cytometry evaluation of cancer stem cell markers ALDH (top), Sca-1$^{-}$CD90$^{+}$ (middle) and Sca-1$^{+}$CD90- (bottom) for PyMT and D2.0R cell lines in 2D (2% FBS tissue-culture treated flasks) as a percentage of parental population (Live, LRC or non-LRC). (**c)** Flow cytometry evaluation of E-cadherin (Ecad) as a percentage of parental population (Live, ALDH$^{+}$, Sca1$^{+}$CD90$^{-}$, and Sca-1$^{-}$CD90$^{+}$) for PyMT and Met-1 cell lines. **(d)** Quantification of sphere formation for each population (ALDH$^{-}$CD90-Sca-1-, ALDH+, ALDH-CD90-Sca-1+, and ALDH-CD90+) for PyMT and Met-1 cell line. **(e)** Quantification of growth of each population (ALDH$^{-}$CD90-Sca-1-, ALDH+, ALDH-CD90-Sca-1+, and ALDH-CD90+) for PyMT, Met-1, and D2.0R cell lines after FACS. P-values were calculated using t test (for normally distributed data) or Wilcoxon test with Bonferroni correction for multiple comparisons when appropriate (* $p < 0.05$, ** $p < 0.01$, *** $p < 0.001$, **** $p<0.0001$).

**Figure S3: ER expression and gating scheme for flow cytometry analysis of CSC populations. (a)** Estrogen receptor alpha expression via immunofluorescence for PyMT, Met-1, D2A1, and D2.0R cell lines. **(b)** Gating scheme for flow cytometry analysis of CSC populations. **(c)** Gating scheme for flow cytometry analysis of CSC populations in the context of cell trace far red (CTFR) label retention.

**Figure S4: Orthotopic tumor growth and initiation is dependent on degree of host immunocompetence.** Tumor volume **(a)** and mass **(b)** for NSG, NODscid, or Balb/c mice injected with 500,000 D2.0R tumor cells in the fourth right and left mammary fat pads. **(c)** Tumor initiation of ALDH+, Bulk, or ALDH- cells in NSG, NODscid, or Balb/c mice.

**Figure S5: Representative BLI images comparing metastasis growth over time for D2.0R-CBG cells in NSG vs NODscid mice.**

**Figure S6: Validation of depletion of NK cells or macrophages. (a)** Schematic for depletion validation experiment in NODscid and Balb/c mice. **(b)** Gating scheme for analysis of NK cells via flow cytometry. **(c**) Quantification of NK cells (% CD49b+ of CD45+ cells) in control mice (NRS) vs Anti-ASGM1 NK depleted mice for Balb/c and NODscid in bone marrow (top) and spleen (bottom). **(d)** Schematic for depletion validation experiment in NODscid mice. **(e)** Gating scheme for analysis of macrophages via flow cytometry. **(f)** Quantification of macrophages (%F4/80+ of CD45+ cells) in bone marrow (left) and spleen (right) for encapsome (control) or clodrosome (macrophage depleted) treated mice.

**Figure S7: NK cell cytotoxicity assays baseline sensitivity for all cell lines tested. (a)** Baseline NK cytotoxicity percentage for Yac-1, PyMT, Met-1, D2.0R, and D2A1 cell lines. **(b**) Colony formation after co-culture with NK cells or without NK cells (control) for D2A1, D2.0R, Met-1, and PyMT cell lines.

**Figure S8: Gating scheme for isolation of quiescent, CTFR+ label retaining cells (LRC) and proliferative, CTFR- non label retaining cells (non-LRC).**

**Figure S9: Gene set enrichment analysis for RNAseq data of quiescent LRC vs proliferative non-LRC demonstrates most activated pathways (up in LRC) are related to inflammation, while those that are suppressed (up in non-LRC) are related to proliferation and metabolism.**

**Figure S10: Knockdown of Bach1 and Sox2 via doxycycline does not alter proliferation or eccentricity. (a)** Number of cells per field of view for D2.0R-CBG cells with doxycycline inducible knockdown of shScr, shBach1, shSox2 and treated with 0, 0.1, 0.5 or 1 ug/mL doxycycline. **(b)** Cell eccentricity as measured by image analysis of D2.0R-CBG cells with doxycycline inducible knockdown of shScr (control), Bach1, or Sox2 and treated with 0, 0.1, 0.5, or 1 ug/mL doxycycline.

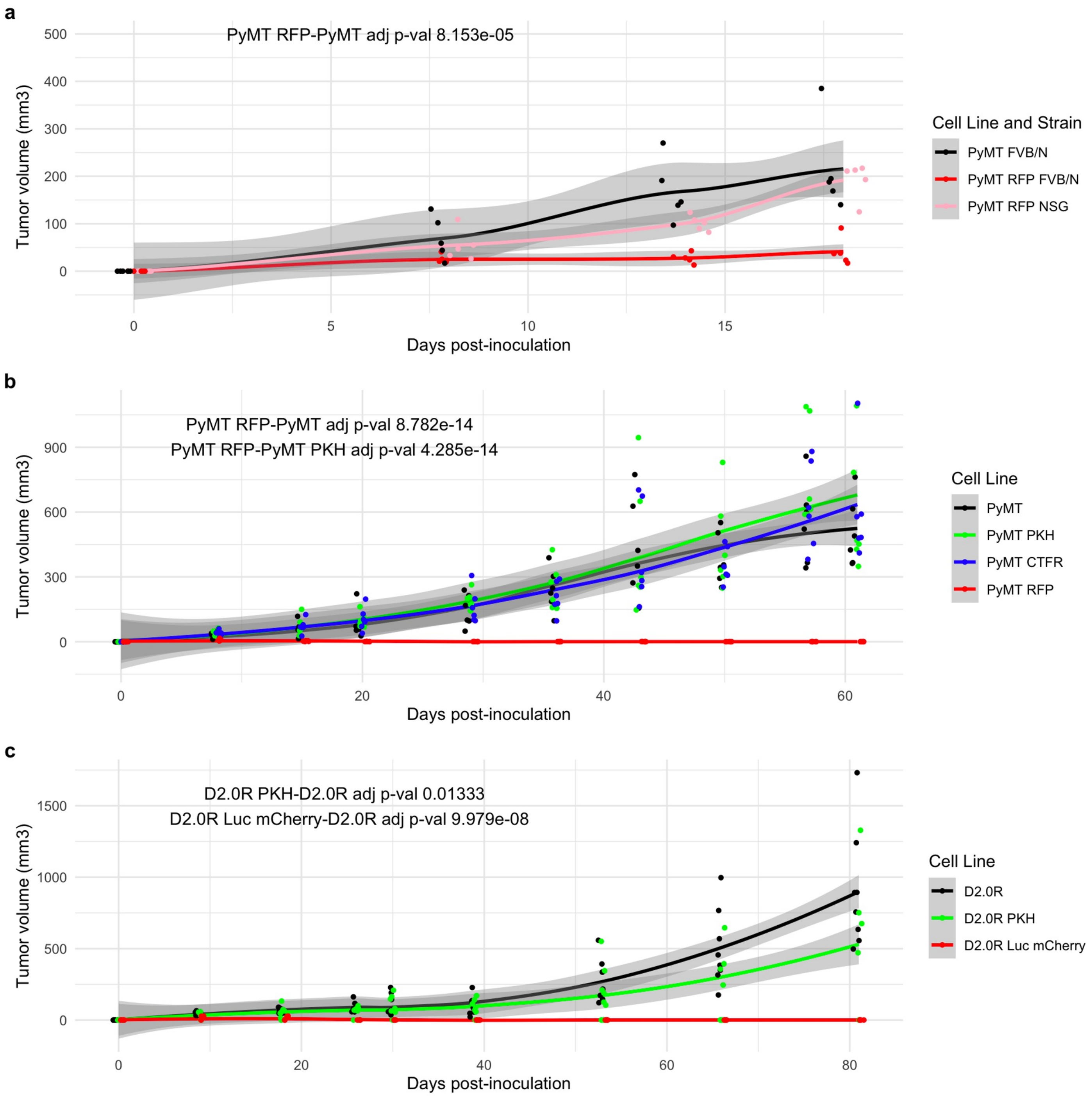
a
PyMT RFP-PyMT adj p-val 8.153e-05
Tumor volume (mm3)
Days post-inoculation
Cell Line and Strain
PyMT FVB/N
PyMT RFP FVB/N
PyMT RFP NSG
b
PyMT RFP-PyMT adj p-val 8.782e-14
PyMT RFP-PyMT PKH adj p-val 4.285e-14
Tumor volume (mm3)
Days post-inoculation
Cell Line
PyMT
PyMT PKH
PyMT CTFR
PyMT RFP
c
D2.0R PKH-D2.0R adj p-val 0.01333
D2.0R Luc mCherry-D2.0R adj p-val 9.979e-08
Tumor volume (mm3)
Days post-inoculation
Cell Line
D2.0R
D2.0R PKH
D2.0R Luc mCherry

Figure S1

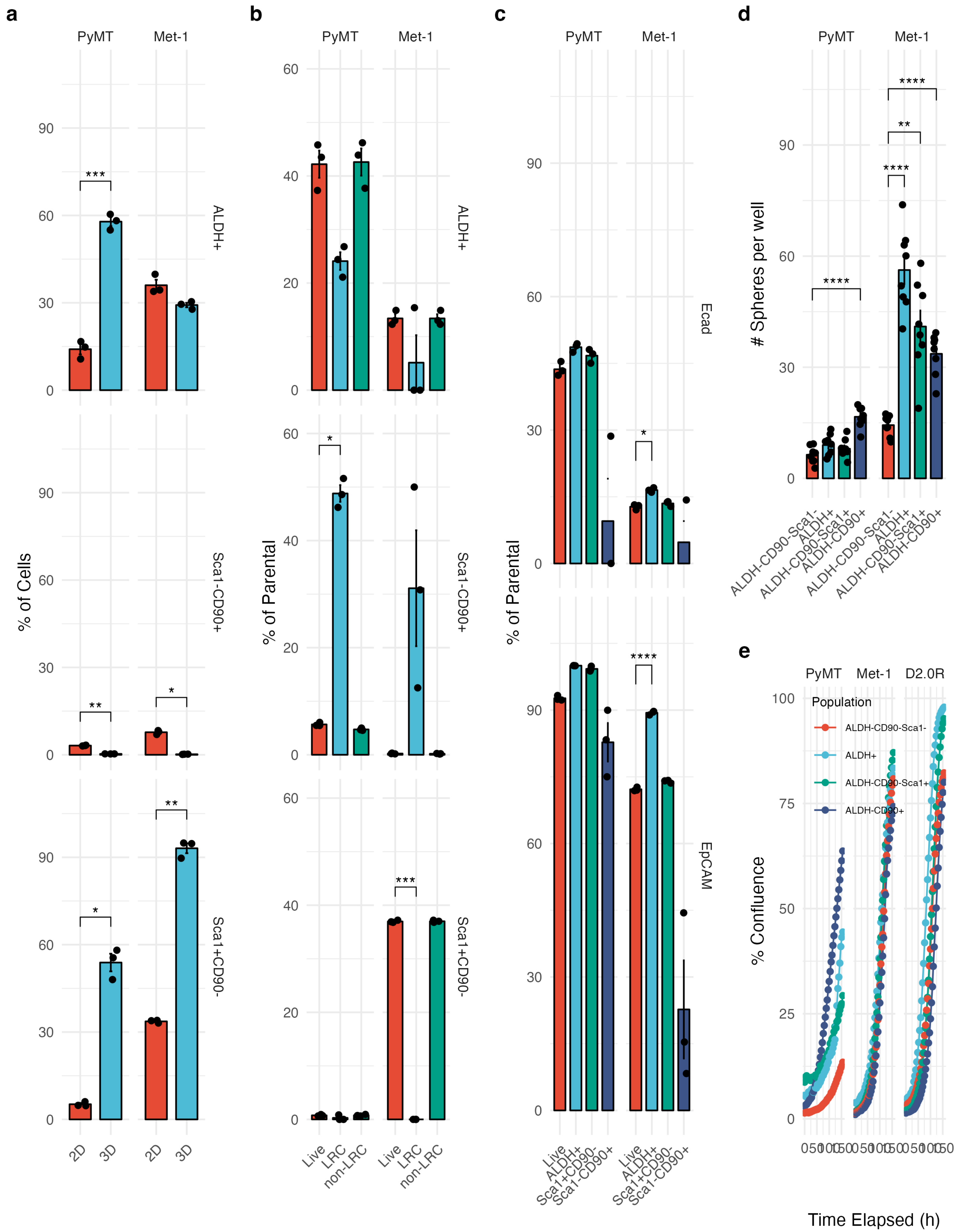
a
b
c
d
e
PyMT
Met-1
ALDH+
Sca1-CD90+
Sca1+CD90-
% of Cells
% of Parental
Ecad
EpCAM
### Spheres per well
% Confluence
Time Elapsed (h)
D2.0R
Population
ALDH-CD90-Sca1-
ALDH+
ALDH-CD90-Sca1+
ALDH-CD90+
2D
3D
Live
LRC
non-LRC
Sca1+CD90-
Sca1-CD90+

Figure S2

a

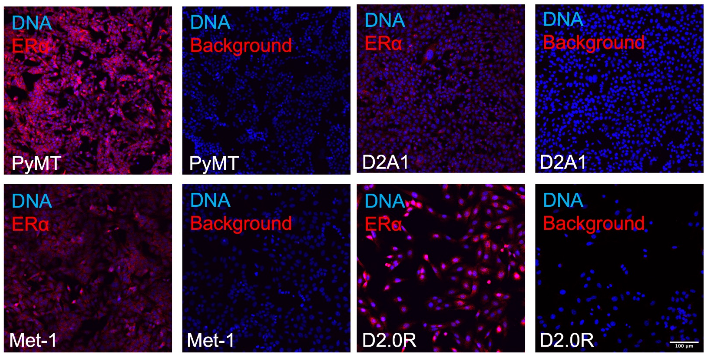


b

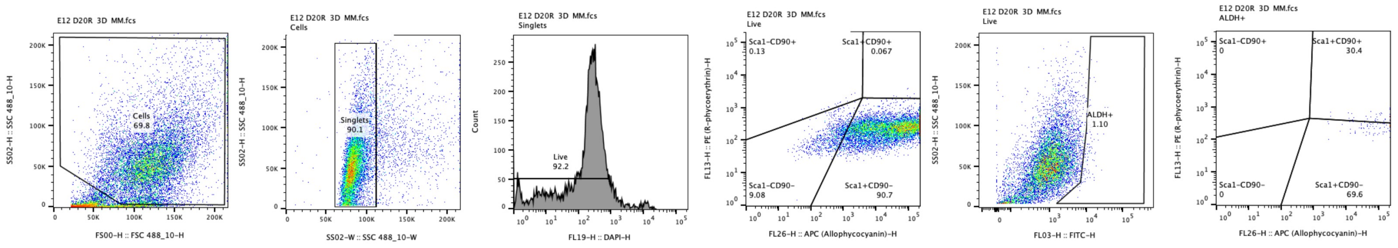


c

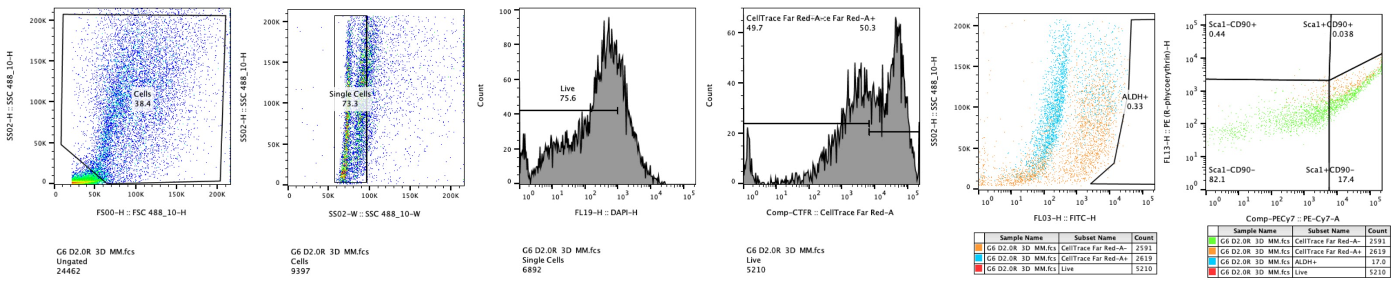


### Figure S3

a

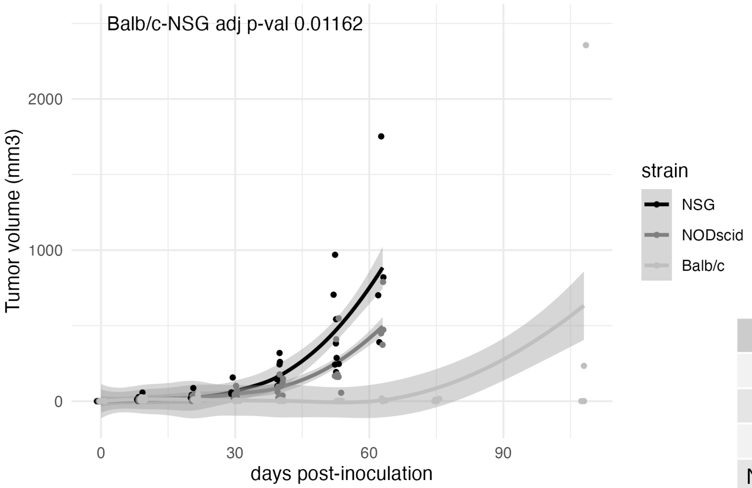


b

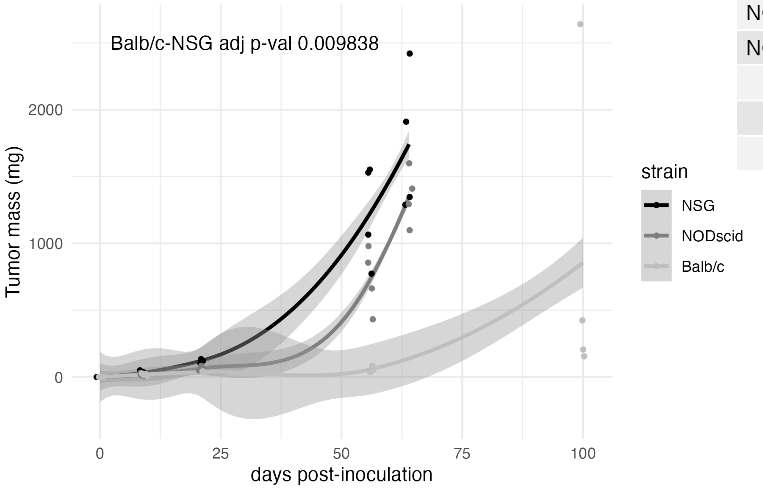


c

| strain | population | 5e5 | 5e4 | 5e3 | TIC frequency |
|---|---|---|---|---|---|
| balb/c | ALDH+ | 1 of 6 | 6 of 6 | 3 of 6 | 1 in 284845 |
| balb/c | Bulk | 5 of 6 | 3 of 6 | 2 of 6 | 1 in 136068 |
| balb/c | ALDH- | 3 of 6 | 3 of 6 | 0 of 6 | 1 in 391315 |
| NODscid | ALDH+ | 6 of 6 | 6 of 6 | 5 of 6 | 1 in 2791 |
| NODscid | Bulk | 6 of 6 | 3 of 6 | 4 of 6 | 1 in 32304 |
| NODscid | ALDH- | 6 of 6 | 6 of 6 | 1 of 6 | 1 in 15351 |
| NSG | ALDH+ | 6 of 6 | 6 of 6 | 5 of 6 | 1 in 2791 |
| NSG | Bulk | 3 of 6 | 5 of 6 | 1 of 6 | 1 in 245525 |
| NSG | ALDH- | 6 of 6 | 4 of 6 | 1 of 6 | 1 in 41571 |

### Figure S4

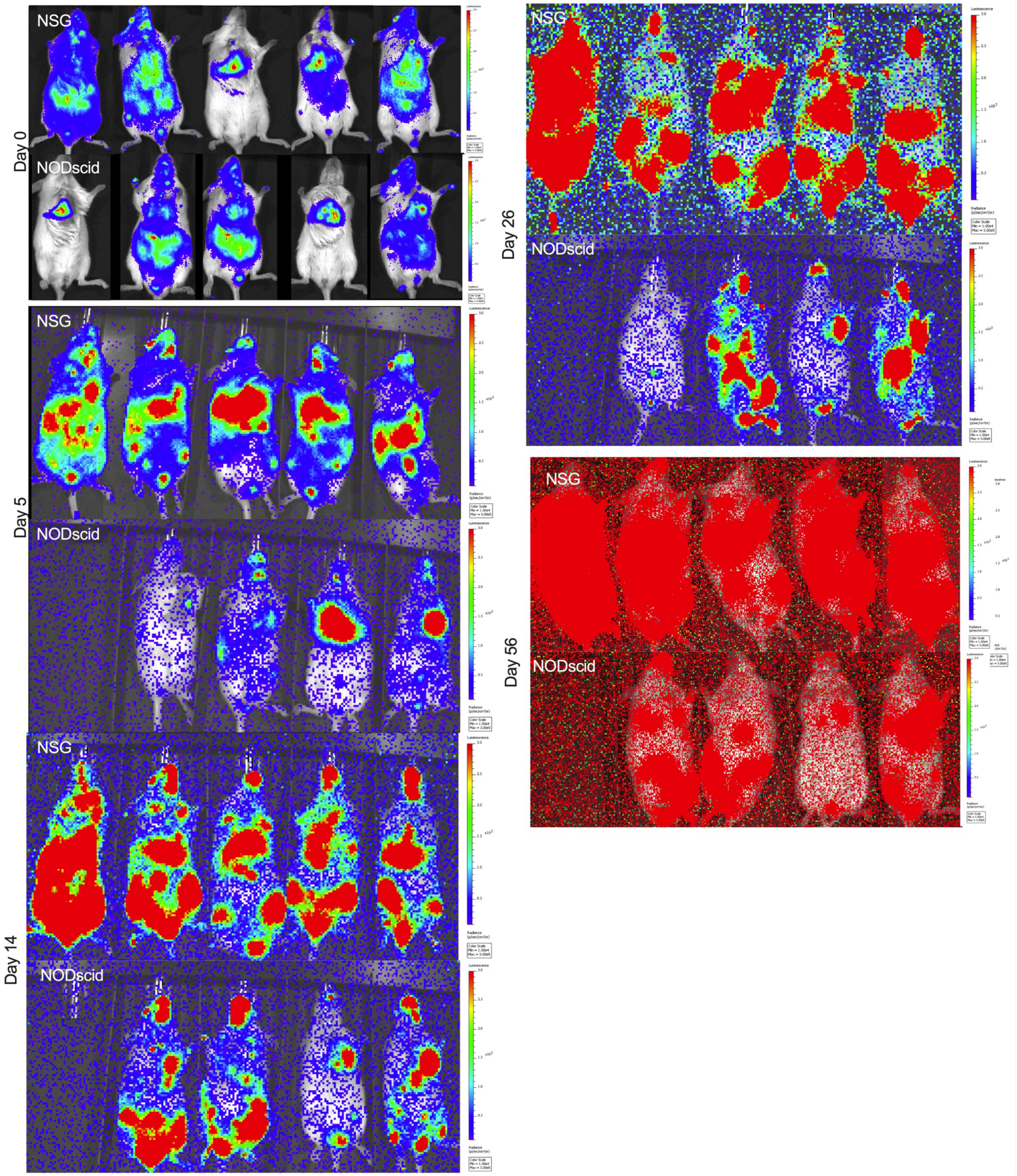
Day 0
NSG
NODscid
Day 5
NSG
NODscid
Day 14
NSG
NODscid
Day 26
NSG
NODscid
Day 56
NSG
NODscid

Figure S5

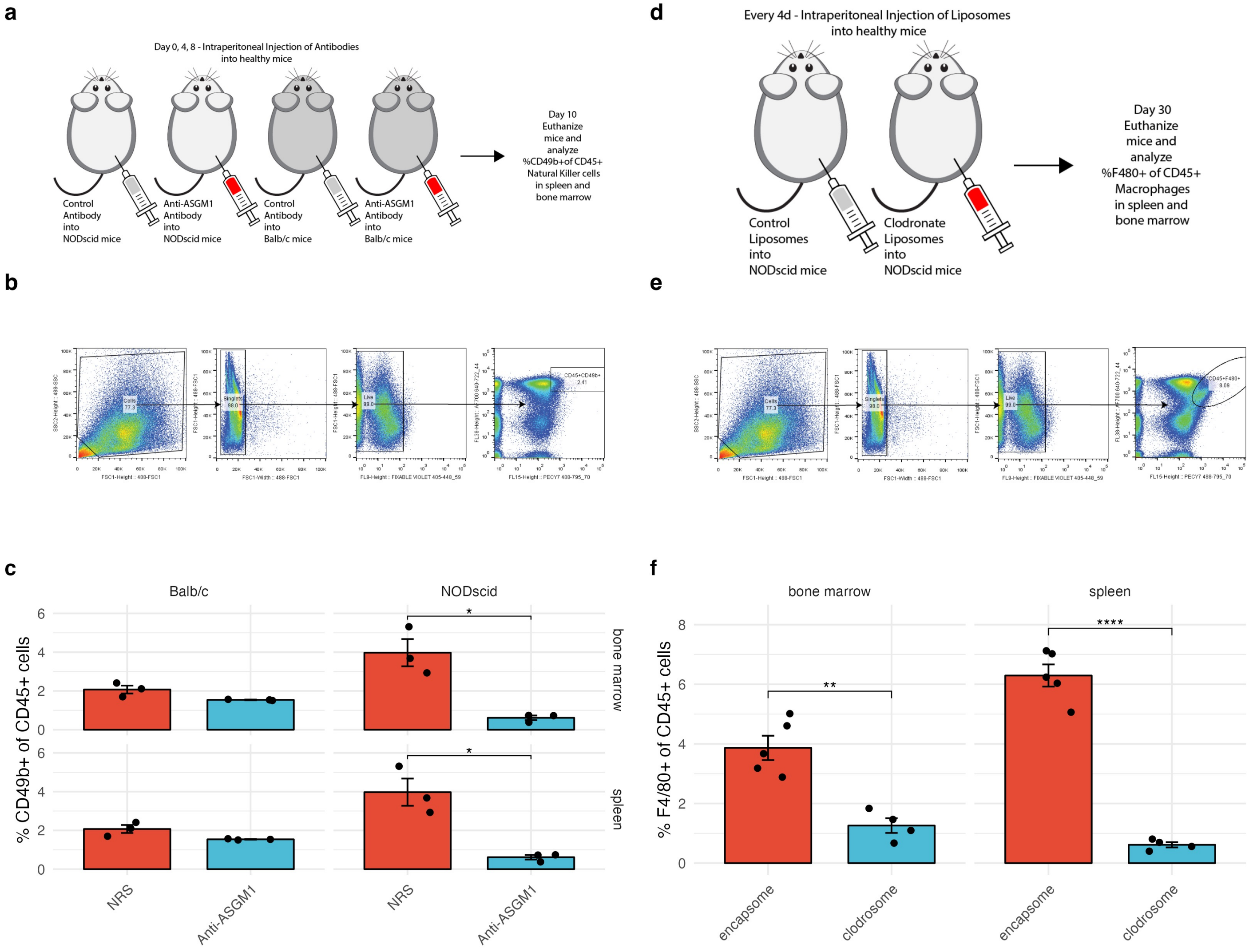
a
Day 0, 4, 8 - Intraperitoneal Injection of Antibodies into healthy mice
Control Antibody into NODscid mice
Anti-ASGM1 Antibody into NODscid mice
Control Antibody into Balb/c mice
Anti-ASGM1 Antibody into Balb/c mice
Day 10 Euthanize mice and analyze %CD49b+of CD45+ Natural Killer cells in spleen and bone marrow
b
Cells 77.3
Singlets 98.0
Live 99.0
CD45+CD49b+ 2.41
c
Balb/c
NODscid
% CD49b+ of CD45+ cells
bone marrow
spleen
NRS
Anti-ASGM1
d
Every 4d - Intraperitoneal Injection of Liposomes into healthy mice
Control Liposomes into NODscid mice
Clodronate Liposomes into NODscid mice
Day 30 Euthanize mice and analyze %F480+ of CD45+ Macrophages in spleen and bone marrow
e
Cells 77.3
Singlets 99.0
Live 99.0
CD45+F480+ 8.09
f
bone marrow
spleen
% F4/80+ of CD45+ cells
encapsome
clodrosome

### Figure S6

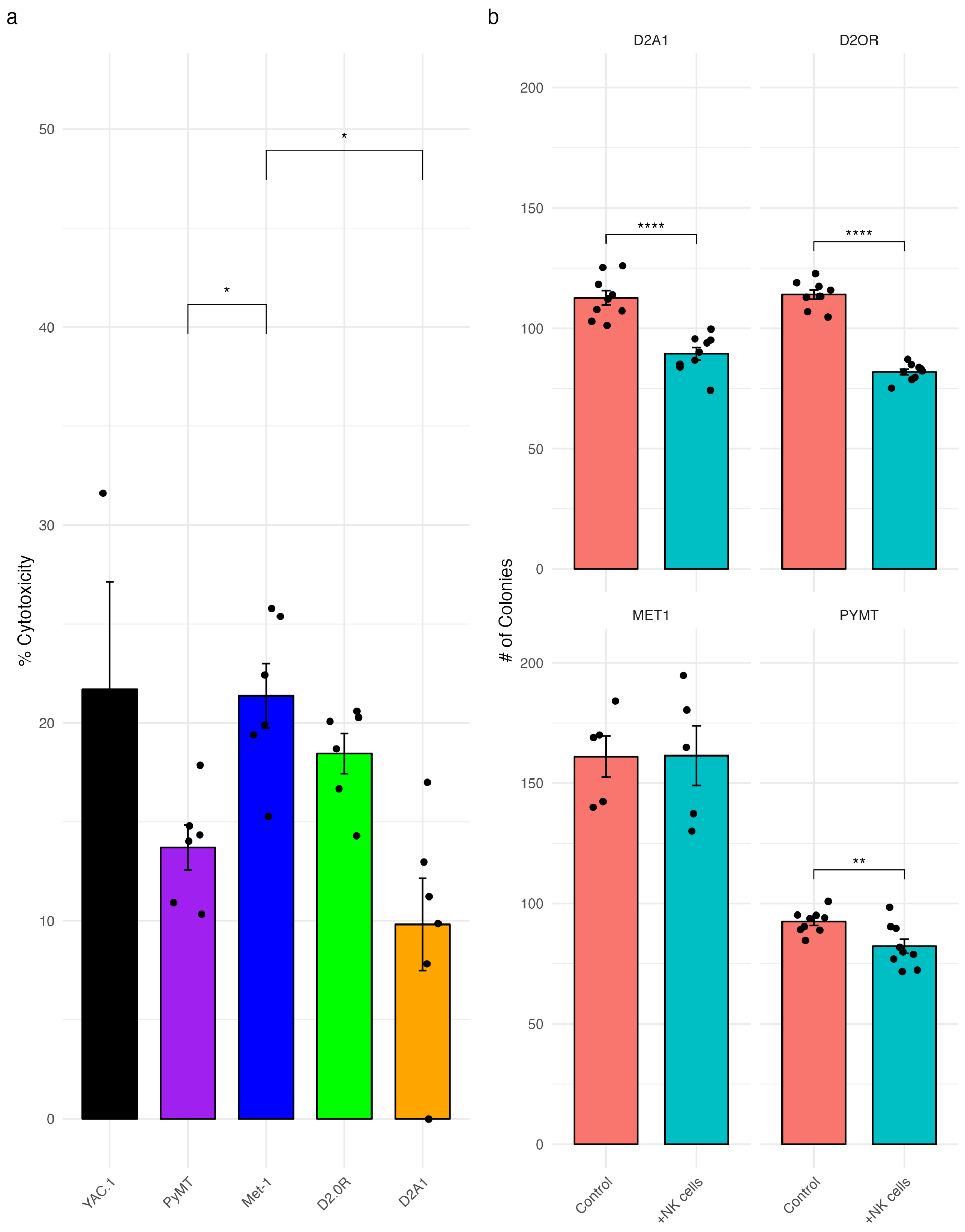
a
50
40
30
20
10
0
% Cytotoxicity
*
*
YAC.1
PyMT
Met-1
D2.0R
D2A1
b
D2A1
D2OR
MET1
PYMT
200
150
100
50
0
### of Colonies
****
****
**
Control
+NK cells

Figure S7

**a**

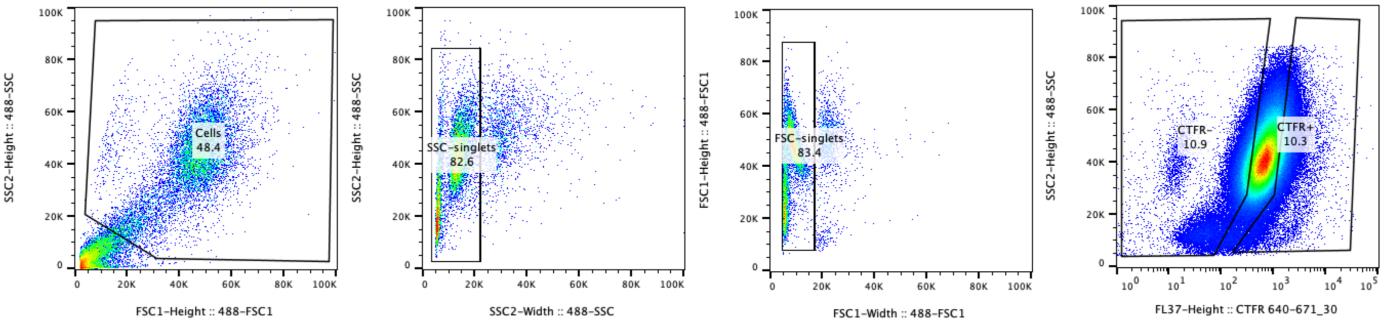

Cells
48.4
SSC2-Height :: 488-SSC
FSC1-Height :: 488-FSC1
SSC-singlets
82.6
SSC2-Width :: 488-SSC
FSC-singlets
83.4
FSC1-Height :: 488-FSC1
FSC1-Width :: 488-FSC1
CTFR-
10.9
CTFR+
10.3
FL37-Height :: CTFR 640-671_30


Figure S8

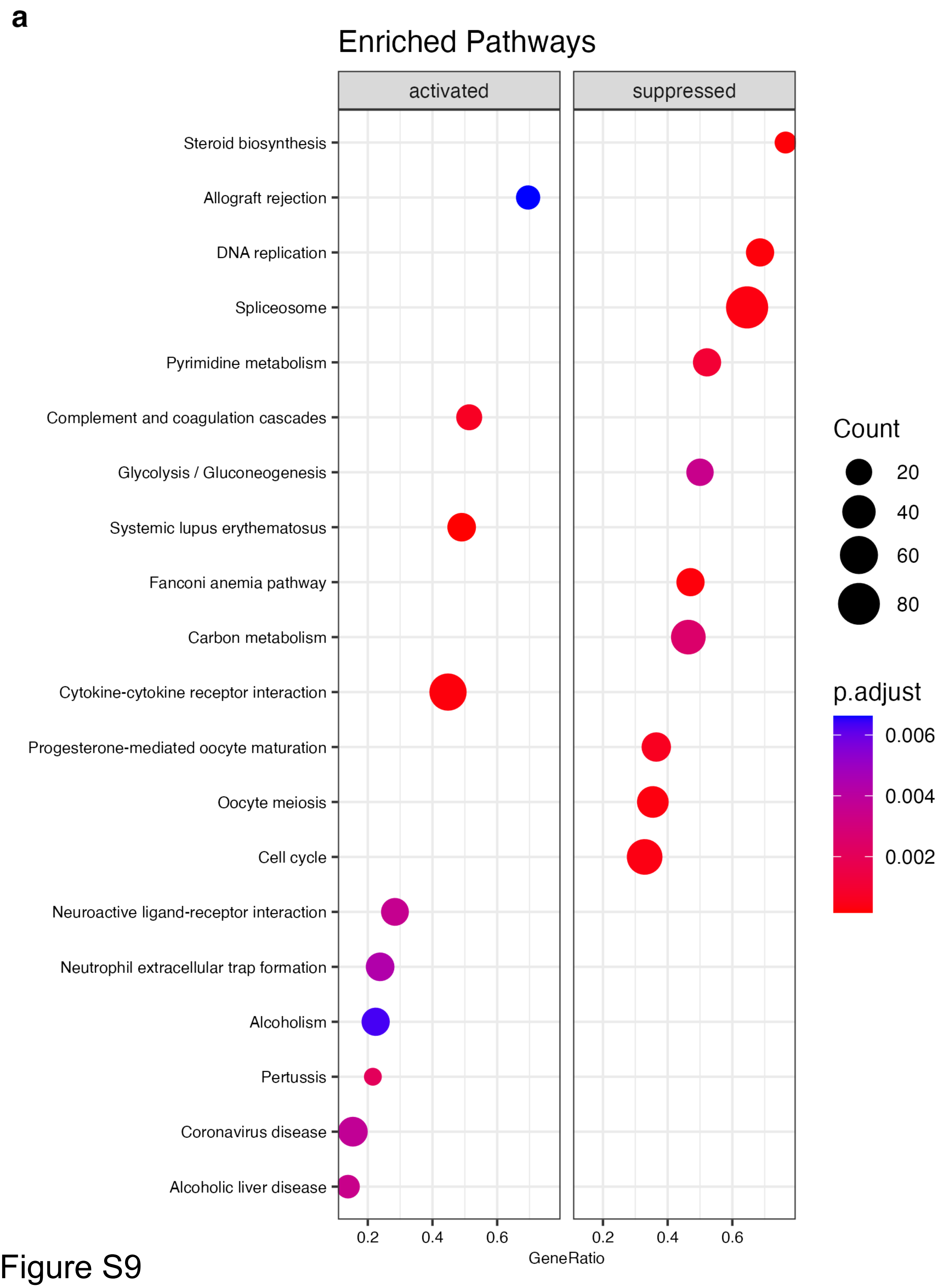
a
Enriched Pathways
activated
suppressed
Steroid biosynthesis
Allograft rejection
DNA replication
Spliceosome
Pyrimidine metabolism
Complement and coagulation cascades
Glycolysis / Gluconeogenesis
Systemic lupus erythematosus
Fanconi anemia pathway
Carbon metabolism
Cytokine-cytokine receptor interaction
Progesterone-mediated oocyte maturation
Oocyte meiosis
Cell cycle
Neuroactive ligand-receptor interaction
Neutrophil extracellular trap formation
Alcoholism
Pertussis
Coronavirus disease
Alcoholic liver disease
0.2
0.4
0.6
0.2
0.4
0.6
GeneRatio
Count
20
40
60
80
p.adjust
0.006
0.004
0.002

Figure S9

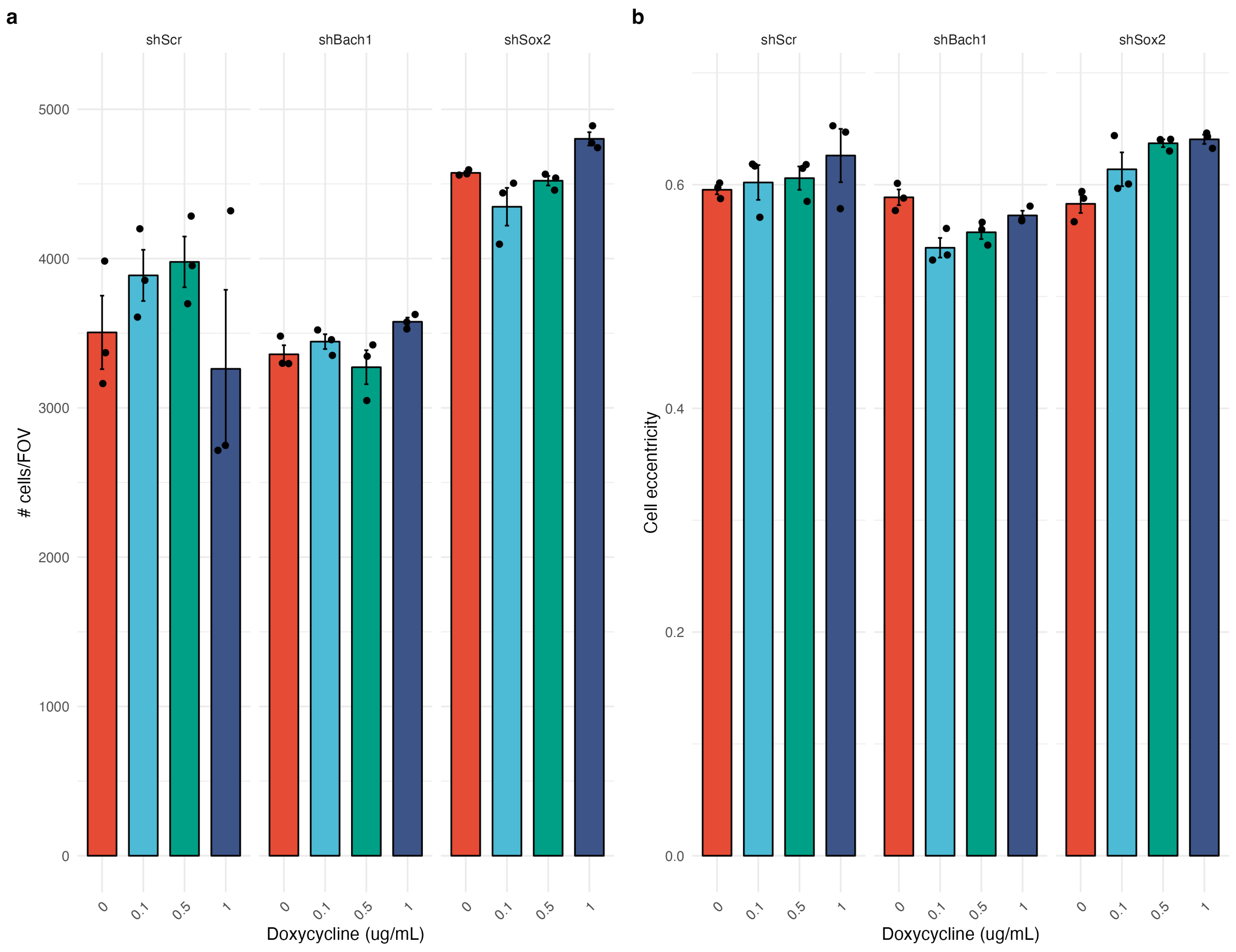

a
shScr
shBach1
shSox2
5000
4000
3000
2000
1000
0
### cells/FOV
0
0.1
0.5
1
Doxycycline (ug/mL)
b
Cell eccentricity
0.6
0.4
0.2
0.0
Doxycycline (ug/mL)


Figure S10
